# Neural modelling of the semantic predictability gain under challenging listening conditions

**DOI:** 10.1101/2019.12.20.872614

**Authors:** Anna Rysop, Lea-Maria Schmitt, Jonas Obleser, Gesa Hartwigsen

## Abstract

When speech intelligibility is reduced, listeners exploit constraints posed by semantic context to facilitate comprehension. The left angular gyrus (AG) has been argued to drive this semantic predictability gain. Taking a network perspective, we ask how the connectivity within language-specific and domain-general networks flexibly adapts to the predictability and intelligibility of speech. During continuous functional magnetic resonance imaging (fMRI), participants repeated sentences, which varied in semantic predictability of the final word and in acoustic intelligibility. At the neural level, highly predictable sentences led to stronger activation of left-hemispheric semantic regions including subregions of the AG (PGa, PGp) and posterior middle temporal gyrus when speech became more intelligible. The behavioural predictability gain of single participants mapped onto the same regions but was complemented by increased activity in frontal and medial regions. Effective connectivity from PGa to PGp increased for more intelligible sentences. In contrast, inhibitory influence from pre-supplementary motor area to left insula was strongest when predictability and intelligibility of sentences were either lowest or highest. This interactive effect was negatively correlated with behavioural predictability gain. Together, these results suggest that successful comprehension in noisy listening conditions relies on an interplay of semantic regions and concurrent inhibition of cognitive control regions when semantic cues are available.

## Introduction

In everyday life, we are remarkably successful in following a conversation even when background noise degrades the speech signal. One important strategy facilitating comprehension in demanding listening scenarios is the prediction of upcoming speech. For example, the sentence fragment “She made the bed with new” provides rich semantic context to inform an accurate prediction of the sentence continuation “sheets”. However, poor semantic context like “We are very pleased with the new” provides little information to build up the prediction of the word “sheets”. This *semantic predictability gain* is a well-documented behavioural phenomenon. It can be observed in healthy young and older listeners (Obleser et al., 2007; Obleser & Kotz, 2010; Vaden et al., 2013, 2015), in hearing-impaired listeners (Holmes et al., 2018), as well as in cochlear implant users (Winn, 2016).

To study the interactive effects of intelligibility and semantic predictability in the listening brain, previous neuroimaging studies used sentences with different occurrence probabilities of the final word while varying the spectral resolution of speech using noise-vocoding (R. V. Shannon et al., 1995). In these studies, the left angular gyrus (AG) has been identified as a key region sensitive to strong semantic constraints in degraded speech (Golestani et al., 2013; Obleser et al., 2007; Obleser & Kotz, 2010). When highly predictable speech was degraded to a moderate extent, activity in AG increased. However, when degradation rendered speech either easily or, if it all, hardly comprehensible, activity in AG dropped irrespective of semantic constraints. Finally, Hartwigsen and colleagues (2015) used focal perturbations induced by transcranial magnetic stimulation to demonstrate that the ventral portion of left AG is functionally relevant for the behavioural predictability gain. Together, these studies provide converging evidence for a role of the left AG in the top-down use of semantic information when comprehension is challenged by degraded speech signals.

The AG is considered a heterogeneous region and has structural connections to language-specific fronto-temporal regions as well as domain-general cingulo-opercular regions (Binder et al., 2009; Seghier, 2013). Based on its cytoarchitectonics, AG can be subdivided into an anterior part (PGa) and a posterior part (PGp; Caspers et al., 2006, 2008).

These subregions have been associated with different functional roles: PGa has been associated with the automatic and domain-general allocation of goal-directed attention and episodic memory. In contrast, PGp was (inconsistently) linked to more controlled and complex semantic-specific processes irrespective of control demands (Bonnici et al., 2016; Bzdok et al., 2016; Jung et al., 2017; Noonan et al., 2013). These findings indicate that the AG might play a central role in language-specific but also in domain-general cognitive operations. Yet, the functional interplay of PGa and PGp within left AG as well as their differential roles in the processing of semantic context in degraded speech are unknown.

As comprehension of degraded speech poses a challenging task on listeners, the brain recruits additional resources beyond the semantic network (Peelle, 2018). These additional resources are thought to be provided by domain-general regions that typically do not show a functional specialization, such as the cingulo-opercular network. The cingulo-opercular network has been implicated in adaptive control processes subserving the flexible allocation of attentional resources and typically comprises the bilateral frontal operculae and adjacent anterior insulae, as well as the dorsal anterior cingulate cortex extending into the pre-supplementary motor area (pre-SMA; Camilleri et al., 2018; Dosenbach et al., 2008; Duncan, 2010). Indeed, previous studies on degraded speech processing have found not only fronto-temporo-parietal regions of the semantic network (Adank, 2012; Alain et al., 2018) but also regions of the cingulo-opercular network (Erb et al., 2013; Vaden et al., 2015, 2017). Additionally, using a word recognition task under challenging listening conditions, Vaden and colleagues (2013) demonstrated that the magnitude of activation within the cingulo-opercular network predicted the word recognition success in the following trial. This upregulation might reflect the activation of additional cognitive resources when no external linguistic cues are available (Peelle, 2018). In contrast, the recruitment of the semantic network is thought to improve comprehension most effectively when semantic context is available. Here, we aim to identify network-level descriptors of the semantic predictability gain to comprehension and investigate how changing task demands shape the adaptive interplay of language-specific and domain-general resources in successful speech comprehension.

Previous studies manipulating predictability and intelligibility of speech suffered from small samples and mainly relied on few levels of noise fixed across participants. Moreover, most of these studies used sparse temporal sampling, precluding the investigation of effective connectivity. Consequently, it remains unclear how the identified regions influence each other during successful comprehension of speech in noise. We aimed to overcome some of the previous shortcomings by performing a continuous event-related functional magnetic resonance imaging (fMRI) experiment with six individualized, varying levels of intelligibility, ranging from largely unintelligible to easily intelligible. Thereby, we accounted for individual differences in auditory perception and assured comparable task performance across participants.

Based on previous studies, we expected a facilitatory behavioural effect of semantic predictability on speech comprehension, which should be strongest at intermediate levels of intelligibility (i.e. when the auditory signal is somewhat degraded but still intelligible) and least pronounced for unintelligible and intelligible sentences. We hypothesized to find increased activation in left AG and other semantic regions as a neural correlate of the semantic predictability gain at intermediate levels of intelligibility. This interaction should be driven by increased effective connectivity between the AG and other semantic regions. Moreover, we expected increasing activity and within-network connectivity in the cingulo-opercular network to improve comprehension in challenging listening conditions (i.e., intermediate intelligibility and low predictability).

As a main finding of our study, highly predictable sentences at intermediate intelligibility levels drove activity in core regions of the semantic network and the behavioural comprehension gain was correlated with higher activity in extended regions of the semantic network. For low predictable sentences, the inhibitory influence between regions within the cingulo-opercular network was less pronounced with increasing intelligibility. In contrast, higher predictability led to stronger inhibitory connectivity within this network when intelligibility increased. The individual degree of the inhibitory influence of predictability and intelligibility on the connection from the pre-SMA to the left insula predicted the individual predictability gain.

## Materials and Methods

### Participants

Thirty healthy young German native speakers took part in this study. After excluding three participants due to excessive head movement during scanning (i.e. range of motion > 1.5 times the voxel size; see Supplementary Figure 1 for an illustration of head movements) and one participant due to reduced task-related activity across the whole brain in the global *F*-test, our final sample yielded 26 participants (age range: 19–29 years, *M* = 25 years; 15 females). All participants were right-handed (mean lateralization index > 90; Oldfield, 1971) and reported no history of neurological or psychiatric disorders as well as no hearing difficulties or disorders.

Participants gave written informed consent prior to participation and received reimbursement of €10 per hour of testing. The study was performed according to the guidelines of the Declaration of Helsinki and approved by the local ethics committee at the Medical Faculty of the University of Leipzig.

### Experimental Procedures

Participants took part in one fMRI session. During this session, we first assessed the individual ability of participants to comprehend speech in noise by tracking the speech reception threshold (SRT). Second, participants performed a repetition task on sentences varying in intelligibility and predictability (Figure 1). The task had a 2 x 6 full factorial design (predictability: high vs. low; intelligibility: -9, -4, -1, +1, +4, +9 dB sentence intensity relative to the individual SRT) with 12 experimental conditions and 18 trials per condition.

**Figure 1.**
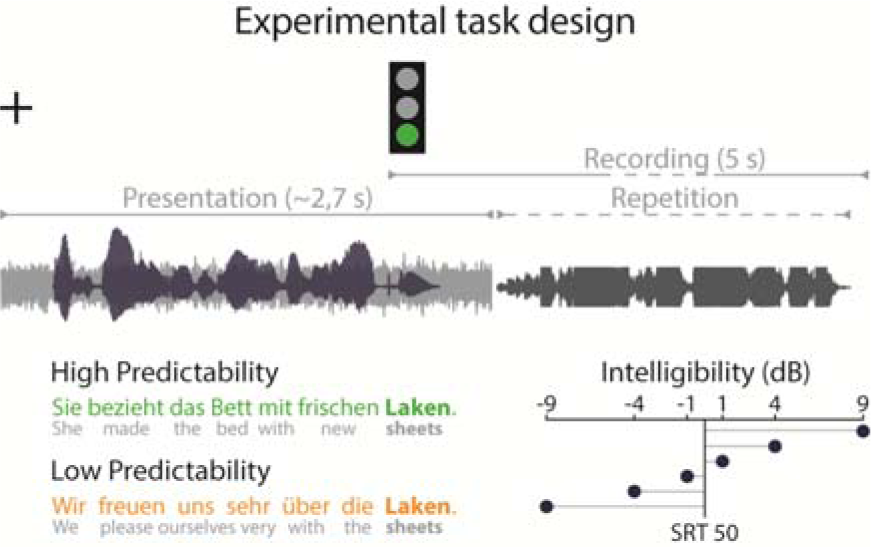
Design of the sentence repetition task. In each trial, participants listened to a sentence (black waveform) embedded in distracting background noise (grey waveform) while viewing a fixation cross on the screen. With the onset of the sentence’s final keyword, a green traffic light prompted participants to orally repeat the whole sentence as accurately as possible within a 5 sec recording period. The preceding sentence context was either predictive (green) or non-predictive of the keyword (orange). Sentences varied orthogonally in intelligibility (ratio of speech intensity to SRT).

#### Adaptive tracking procedure of the speech reception threshold

To account for inter-individual differences in speech perception acuity in the main experiment, we determined each participant’s SRT beforehand, that is, the signal-to-noise ratio (SNR) required to correctly repeat a sentence with a probability of 50%. In an adaptive up-down staircase procedure (Kollmeier et al., 1988), participants listened to 20 sentences with highly predictable final words energetically masked by speech-shaped noise. After the presentation of each sentence, participants repeated the whole sentence as accurately as possible and the investigator directly rated the response as either correct or incorrect. We rated a response as incorrect if any word in the repeated sentence was missing, incomplete or inaccurately inflected. A correct repetition was followed by an SNR decrease (i.e. the following sentence was less intelligible), an incorrect repetition was followed by an SNR increase (i.e. the following sentence was more intelligible). The tracking procedure started with an initial SNR of 5 dB and a trial-to-trial step size of 6 dB. A turning point (i.e., an increase in SNR is followed by a decrease or vice versa) reduced the step size by a factor of 0.85. We determined SRTs as the average of SNR levels presented on those trials leading to the final five turn points. SRTs had an average of +1.7 dB SNR (SD = 2.3, range: -2.1 to +6.4 dB). All sentences were presented in the MR scanner while running the fMRI sequence later used in the experiment. Individual SRTs were used as a reference for the manipulation of intelligibility in the main experiment to account for inter-individual differences in speech-in-noise comprehension and thereby effectively modulate behavioural performance.

#### Stimulus material

To manipulate predictability of speech in the sentence repetition task, we used the German version of the speech in noise (SPIN) corpus (Kalikow et al., 1977; for a detailed description of the German version, see Erb et al., 2012). The German SPIN corpus contains pairs of spoken sentences which have the same final word (i.e. keyword) but differ in cloze probability of the keyword (i.e. the expectancy of a word given the preceding sentence; Taylor, 1953). One sentence of a pair provides poor semantic context for the prediction of the keyword (“We are very pleased with the new <u>sheets</u>.”; low predictability), whereas the complementary sentence provides rich context for the same keyword (“She made the bed with new <u>sheets</u>.”; high predictability). More specifically, the predictability of the keyword varies with the number of pointer words in a sentence that semantically link to the sentence-final word (e.g., “made”, “bed”, and “new” in the above example). Keywords from sentences with rich context had a mean cloze probability of 0.85 (*SD* = 0.14), low predictable keywords had a mean cloze probability of 0.1 (*SD* = 0.02).

In addition to the 100 original sentence pairs, we extended the corpus by another 36 sentences which were constructed and recorded during the development of the original German SPIN corpus. Initially, these sentences were not included in the corpus, as a smaller set of sentences was sufficient for the purposes of the developers. To obtain predictability ratings for the keywords in the new sentences, 10 participants who did not take part in the fMRI experiment (age range: 24–28 years; *M* = 26 years; 4 females) performed a written sentence completion test (low predictability: < 5 participants reported the correct keyword; high predictability: > 5 participants). Together, 8 new sentence pairs and the original German SPIN sentence pairs yielded 216 experimental sentences (*M* = 2143 ms, *SD* = 256 ms). The remaining 20 new sentences were used in the adaptive tracking procedure prior to the experiment (see Supplementary Figure 2 for a detailed illustration of how sentences were assigned to the experiment and the adaptive tracking procedure).

Intelligibility of speech was manipulated by varying the intensity of the experimental sentences relative to a constant signal intensity of speech-shaped noise in the background. We created spectrally speech-shaped noise by filtering white noise with the long-term average spectrum of all experimental sentences (e.g. Nilsson et al., 1994). The noise stream preceded and ensued single sentences by 250 ms. Our six experimental intelligibility levels were symmetrically spaced around individual SRTs and logarithmically increased towards the extremes. With our intelligibility levels, we aimed to cover the full range of behavioural performance in each participant and sample intermediate intelligibility in smaller steps as SNR changes in this range affect performance more strongly.

#### Sentence repetition task

Participants performed an overt sentence repetition task during fMRI. They listened to single sentences while viewing a white fixation cross on a black screen. With the onset of the keyword, a green traffic light appeared on the screen and indicated the start of a 5000 ms recording period. During the recording period, participants orally repeated the sentence as accurately as possible. Whenever participants did not comprehend the complete sentence, they repeated the sentence fragments they could grasp. If participants could not repeat any word, they said that they did not understand the sentence. The intertrial interval was randomly set between 2000 and 7000 ms and served as an implicit baseline for the fMRI analysis. Sentences were presented in a pseudorandomized order avoiding the repetition of one intelligibility level more than three times in a row to prevent adaptation to specific intelligibility levels. The experiment was split in six blocks of approximately 8 minutes which were intermitted by pauses of 20 s each.

All sentences were presented at a comfortable volume. Sound was played via MR-Confon headphones (Magdeburg, Germany) and recorded via a FOMRI-III microphone (Optoacoustics, Yehuda, Israel). Stimulus presentation was controlled by Presentation software (version 18.0, Neurobehavioral Systems, Berkeley, USA, www.neurobs.com). The experiment lasted 50 minutes.

#### MRI acquisition

Whole brain fMRI data were acquired in one continuous run with a 3 Tesla Siemens Prisma Scanner and 32-channel head coil, using a dual gradient-echo planar imaging multiband sequence (Feinberg et al., 2010) with the following scanning parameters: TR = 2,000 ms; TE = 12 ms, 33 ms; flip angle = 90°; voxel size = 2.5 x 2.5 x 2.5 mm with an interslice gap of 0.25 mm; FOV = 204 mm; multiband acceleration factor = 2. In sum, 1,500 volumes were acquired per participant, each consisting of 60 slices in axial direction and interleaved order. To increase coverage of anterior temporal lobe regions, slices were tilted by 10° off the AC-PC line. Field maps were acquired for later distortion correction (TR = 620 ms; TE = 4 ms, 6.46 ms). Additionally, high-resolution T1-weighted images were either obtained from the in-house database if available or were acquired prior to the functional scans with an MPRAGE sequence (whole brain coverage, TR = 1300 ms, TE = 2.98 ms, voxel size = 1 x 1 x 1 mm, matrix size = 256 x 240 mm, flip angle = 9°).

### Data Analysis

#### Behavioural Data Analysis

All spoken response recordings from the main experiment were cleaned from scanner noise using the noise reduction function in Audacity (version 2.2.2, https://www.audacityteam.org/). For transcriptions, we split the recordings in two independent sets and assigned each recording set to one of two raters. To validate the quality of transcripts, a third rater additionally transcribed half of the recordings in each set. Cohen’s kappa coefficient (Cohen, 1960) indicated very good inter-rater reliability for keywords (rater 1 and 3:κ = .94; rater 2 and 3:κ = .97). Finally, one rater manually determined onset and duration of oral responses.

As only predictability of the final keyword but not its preceding context is explicitly manipulated in the German SPIN corpus, we limited behavioural analyses to keywords. After rating keywords as incorrect (including missing, incomplete and inaccurately inflected keywords) or correct, we calculated the proportion of correctly repeated keywords for each participant and condition.

In the behavioural analysis, we quantified the effects of predictability and intelligibility on speech comprehension. For each participant, we fitted psychometric curves to the proportion of correct keywords across intelligibility levels, separately for sentences with low and high predictability. In detail, we fitted cumulative Gaussian sigmoid functions as beta-binomial observer models using the Psignifit toolbox (Fründ et al., 2011) in MATLAB (version R2018b, MathWorks). The beta-binomial model extends the standard binomial model by an overdispersion parameter which allows to carry out statistical inference on data affected by performance fluctuations challenging the serial independence of trials (Schütt et al., 2016). To estimate all five parameters describing the function (guess and lapse rate, threshold, width and overdispersion), we applied Psignifit’s default priors for experiments with binary single-trial outcome (here: correctly vs. incorrectly repeated keyword; Schütt et al., 2016).

Goodness of psychometric fits was assessed by comparing the empirical deviance (i.e., two times the log-likelihood ratio of the saturated model to the fitted model) with a Monte Carlo simulated deviance distribution (Wichmann & Hill, 2001). For each psychometric curve, a reference distribution was created by 1) randomly drawing from a binomial distribution with *n* = 18 trials and *p* = fitted probability of a correct response for each intelligibility level, 2) calculating the deviance across intelligibility levels, and 3) repeating this procedure for 10,000 samples. All empirical deviances fell within the 97.5% confidence interval of their respective reference distribution, indicating that psychometric curves properly represented observed behavioural data in every participant. There was no significant difference between the deviance of fits for sentences with low vs. high predictability (*t*_25_ = 0.21, *p* = 0.839, *r* = 0.21, BF_10_ = 0.21). Parameter estimates including credible intervals for single participants can be found in Supplementary Figure 3. To illustrate goodness of fit, we additionally calculated R^2^ based on the Kullback-Leibler divergence (*R*^2^_*KL*_) which represents the reduction of uncertainty by the fitted model relative to a constant model (Cameron & Windmeijer, 1997). On average, psychometric curves of sentences with high predictability yielded an of 0.96 (*SD* = 0.04, range: 0.84-0.998), sentences with low predictability an (*R*^2^_*KL*_) of 0.94 (*SD* = 0.05, range: 0.79-0.997).

As all 12 experimental conditions become subsumed in two psychometric curves, interaction effects can be evaluated by comparing parameter estimates between curves for sentences with low and high predictability. The threshold represents the intelligibility level at which participants correctly repeat half of the keywords. Keeping all other parameters constant, a threshold decrease indicates a comprehension gain at intermediate but not high and low intelligibility levels. We hypothesized that the interaction effect of intelligibility and predictability manifests in a smaller threshold for sentences with high predictability when compared to low predictability. Additionally, we expected that an increase in intelligibility leads to a stronger comprehension gain for sentences with high relative to low predictability. The sensitivity to changes in intelligibility is reflected by the slope which is inversely related to the width and here describes the steepness of a psychometric curve at a proportion correct of 0.5. To explore whether also other parameters of psychometric curves were modulated by the predictability of sentences, we compared guess and lapse rate (i.e., lower and upper asymptote) as well as width between sentences with high and low predictability. Parameter estimates of the two psychometric curves were contrasted using a paired-sample *t*-test (or a Wilcoxon signed-rank test whenever the assumption of normality was not met). Additionally, we regressed out SRTs from psychometric parameter estimates and reran all *t*-tests between sentences with low and high predictability to control for potential confounding effects of overall inter-individual differences in auditory perception.

We quantified the strength of evidence in favour of our hypotheses by calculating the Bayes Factor (BF; Jeffreys, 1961) for all *t*-tests and correlations carried out in the behavioural analyses using the default settings in JASP (i.e., comparing against a Cauchy prior with scale = √2 / 2; version 0.8.6.0). Whereas BF_10_ = 1 supports neither the alternative nor the null hypothesis, BF_10_ > 3 indicates increasingly substantial support for the alternative hypothesis (Kass & Raftery, 1995). For BF_10_ = 3, the data are 3 times more likely to have occurred under the alternative hypothesis than the null hypothesis. As an additional effect size measure, we report Pearson’s correlation coefficient *r* (Rosenthal & Rubin, 1994).

#### Functional MRI Data

Functional MRI data were analysed using the Statistical Parametric Mapping software (SPM12, version 7219, Wellcome Department of Imaging Neuroscience, London, UK) and MATLAB (version R2017b). Preprocessing steps were applied in line with SPM’s default settings and comprised realignment, distortion correction, segmentation, co-registration to the individual high-resolution structural T1-scan and spatial normalization to the standard template by the Montreal Neurological Institute (MNI) while keeping the original voxel size. Volumes were smoothed using a 5 mm full width at half maximum Gaussian kernel to allow statistical inference based on Gaussian random-field theory (Friston et al., 1994). Realignment parameters were obtained from the first echo. Temporal signal-to-noise ratio maps were calculated for each echo based on the first 30 volumes of each participant. Echoes were combined using the information from the corresponding temporal signal-to-noise ratio maps as weighting factors and realigned to the first scan. This approach has been proposed to increase SNR in brain regions typically suffering from signal loss (e.g., anterior temporal lobe regions; see Halai et al., 2014 for a similar approach). Motion parameters were calculated from the parameters obtained by realignment. Additionally, framewise displacement as described by Power et al., 2012 was calculated as an index of head movement. Volumes that exceeded a framewise displacement of 0.9 (Siegel et al., 2014) were included as nuisance regressors.

At the single-participant level, we set up a general linear model (GLM). For each experimental condition, onset and duration of the sentence presentation period were modelled with a stick function and convolved with SPM’s canonical hemodynamic response function. Additionally, we modelled onset and duration of the sentence repetition periods using individual speech onset times and speech durations. We further included motion parameters obtained from the realignment step and framewise displacement as regressors of no interest. Data were high-pass filtered with a cut-off at 128 s and serial correlations were taken into account using a first-order autoregressive model. Each experimental regressor was contrasted against the implicit baseline (intertrial interval), thus yielding 12 contrast images of interest, one for each experimental condition. Additionally, an *F*-contrast containing the experimental regressors was set up to capture experimental effects of interest and exclude effects of no interest (i.e. sentence repetition periods, motion-related activity).

At the group level, these contrasts were submitted to a random-effects flexible factorial design with the experimental factors predictability (high, low) and intelligibility (-9 to +9 dB) entered as interaction term, and an additional factor modelling the subjects’ constant. First, we set up a *t*-contrast modelling a linear increase in intelligibility (-9 > -4 > -1 > +1 > +4 > +9 dB SNR) to reveal brain regions tuning to increasing clarity of speech irrespective of predictability (i.e. main effect of intelligibility).

Further, we were interested in those brain regions showing a linear or quadratic interaction of intelligibility and predictability. The interaction contrasts were set up as separate t-contrasts for linearly (high > low, -9 > -4 > -1 > +1 > +4 > +9 dB SNR relative to SRT) and quadratically modelled intelligibility (high > low, -9 < -4 < -1 = +1 > +4 > +9 dB). These parametric interaction contrasts were set up in both directions (high > low / low > high).

Single-participant’s parameter estimates were extracted from significant peak voxels and transformed into percent signal change using the rfx toolbox (Gläscher, 2009) for illustration purposes. Figures were created using MRIcroGL (www.mricro.com, version v1.0.20180623).

All effects for the whole-brain group analysis are reported at a voxel-level family-wise (FWE) correcting threshold of *p* < 0.05 to control for multiple comparisons. We do not report clusters with less than 10 voxels, as we consider such small clusters biologically implausible. Local maxima are reported in MNI space. We used the SPM Anatomy Toolbox (Eickhoff et al., 2005, version 2.2c) and the Harvard-Oxford cortical structural atlas (https://fsl.fmrib.ox.ac.uk/fsl/fslwiki) to classify the anatomical correspondence of significant voxels and clusters.

#### Brain–Behaviour Correlation

We identified brain regions reflecting the behavioural predictability gain across intelligibility levels by correlating neural response patterns with individual task performance. On the behavioural level, we extracted the proportion of correctly repeated keywords in each experimental condition and participant as modelled by psychometric curves.

To quantify how strongly the proportion correct in each intelligibility level was affected by the predictability of sentences for single participants, we weighted the interaction contrast by behaviour. The interaction contrast was built from the Kronecker product of differential predictability and intelligibility effects. A differential effect is the transposed orthonormal basis of differences between columns of an *l*-by-*l* identity matrix (predictability: *l* = 2, intelligibility: *l* = 6; for a detailed description see Henson, 2015). Multiplying this F-contrast with the condition-wise task performance of each participant yielded the proportion of variance in performance explained by the interaction, i.e. the predictability gain (or loss) in each intelligibility level.

On the neural level, we limited our analysis to all voxels consistently implicated in task-related activity as indicated by a significant *F*-contrast (*p*_uncorrected_ < 0.05) calculated across single-participant parameter estimates of all experimental conditions (see mask in Supplementary Figure 4). For each voxel within the mask, we correlated single-participant behavioural predictability gain and neural parameter estimates across conditions, using a custom Matlab script. After applying Fisher’s z-transformation to all Pearson product-moment correlation coefficients, single-participant maps were submitted to the Local Indicators of Spatial Association (LISA) group-level one-sample *t*-test (Lohmann et al., 2018). LISA is a threshold-free framework that allows to find consistent effects in small brain regions by applying a non-linear filter to statistical maps before controlling for multiple comparisons at a false discovery rate (FDR) < 0.05. Voxel-wise FDR scores were obtained from a Bayesian two-component mixture model in which filtered statistical values are compared to a null distribution based on 5000 permutations (i.e., randomly switching signs of statistical maps in participants).

### Dynamic Causal Modelling (DCM)

To investigate condition-specific interactions between brain areas, we conducted two separate bilinear one-state deterministic Dynamic Causal Modelling (DCM) analyses using the DCM 12.5 implementation in SPM 12. DCM is a forward-model-based method that models the dynamics of hidden neuronal states within a predefined set of regions and relates them to the measured blood-oxygen-level-dependent (BOLD) signal (Friston et al., 2003). Using this Bayesian framework, modulations of effective connectivity by experimental conditions can be assessed.

#### Seed region selection

According to our main hypothesis, we were interested in the neural network dynamics underlying the predictability gain in speech in noise processing and the role of left angular gyrus. In the GLM analysis, we found two distinct networks implicated in the neural interaction of predictability and intelligibility: the semantic network and the cingulo-opercular network (CO). Here, we test effective connectivity in a subset of regions within each network. The semantic model was based on the GLM interaction contrast (high > low, linearly modelled intelligibility) and included left posterior middle temporal gyrus (pMTG), PGa and PGp as seed regions. This model specification allowed to investigate differential contributions of subregions within AG, namely PGa and PGp. Based on the opposite GLM interaction contrast (low > high, linearly modelled intelligibility), we specified the CO model including pre-SMA/paracingulate gyrus as well as left and right anterior insula as seed regions.

For each subject, we identified the peak voxel nearest to the group maximum of each seed region within a sphere of 10 mm radius using the appropriate first-level contrast. Individual timeseries of each region (summarized as the first eigenvariate) were extracted from all voxels within a sphere of 6 mm radius centered on the individual maximum and exceeding the liberal threshold of *p*_uncorrected_ < 0.05. The extracted timeseries were adjusted to the respective effects-of-interest *F*-contrast to exclude variance of no interest. The exact anatomical localization of individual seed regions was verified using SPM’s Anatomy Toolbox (Eickhoff et al., 2005). For spatially adjacent PGa and PGp, we made sure that seed regions were not overlapping within each subject.

The summarized timeseries together with information about the onsets of the relevant conditions served as input for the DCM. The design matrix for the DCM analyses differed from the design matrix used in the GLM analysis in the following way: seven regressors were defined, modelling the onsets of (1) all experimental stimuli, (2) high predictable sentences at low intelligibility levels (-9, -4 dB SNR relative to SRT), (3) high predictable sentences at medium intelligibility levels (-1, +1 dB SNR relative to SRT), (4) high predictable sentences at high intelligibility levels (+4, +9 dB SNR relative to SRT). The remaining regressors modelled the onsets of low predictable sentences at low, medium and high intelligibility, respectively. The first regressor served as sensory input (i.e. driving input) of the stimuli into the model. Regressors (2) to (7) were used as modulatory inputs and encoded predictability and intelligibility of the stimulus. Considering that each experimental condition comprised only 18 trials, we aimed to increase the signal-to-noise ratio of our DCM parameter estimates by binning neighbouring intelligibility levels into groups of low, medium and high intelligibility. As inputs were not mean-centered, intrinsic parameter estimates can be interpreted as average connection strength in the absence of experimental manipulations (Marreiros et al., 2008).

#### DCM model architecture

At the first-level, 84 models were specified for each subject. All models consisted of three types of parameters: (i) intrinsic connections between two nodes (unidirectional or bidirectional), (ii) changes in coupling strength between regions dependent on the experimental manipulation (modulatory effects) and (iii) direct influence of external input on a region (driving input).

For both DCM analyses, we modelled full reciprocal intrinsic connections as well as (inhibitory) self-connections. The driving input was set to either one region at a time, or all possible combinations between regions (resulting in 7 possible combinations). The following potential modulatory influences were considered: either one connection (except for the self-connections) was modulated separately or two afferent or efferent connections were modulated by all experimental conditions at a time (12 possible combinations; see Supplementary Figure 5 for a schematic overview of the model space). Since family-level inference has been found especially useful in case of large model spaces (Penny et al., 2010), the resulting model space of 7 x 12 models was partitioned into 12 families, grouping models according to the modulatory input (i.e. within a family, the modulating connection was kept constant while the driving input was varied). We used the same model architecture for both DCM analyses.

All 84 models per participant were inverted using a Variational Bayes approach to estimate parameters that provide the best trade-off between accuracy and complexity quantified by free energy (Friston et al., 2006, 2007). At the second level, a random-effects Bayesian Model Selection procedure was applied to identify the most probable family of models given the fMRI data quantified by the respective exceedance probability (Penny et al., 2010; Stephan et al., 2009). Single-subject parameter estimates were averaged across all models within the winning family using Bayesian Model Averaging. Intrinsic parameter estimates were passed to a one-sample *t*-test. Modulatory parameter estimates were submitted to a two-way repeated measures analysis of variance (ANOVA) with the within-subject factors predictability (high, low) and intelligibility (low, medium, high) to investigate the interaction effect on modulated connections between regions. ANOVAs were calculated in JASP (version 0.9.1); Greenhouse-Geisser correction was applied if Mauchly’s test indicated that the assumption of sphericity was not met. The strength of ANOVA effects was quantified with partial eta squared 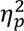.

Finally, we investigated the within-subject relationship of behavioural comprehension and functional connectivity in the brain. For single participants, we resorted to the condition-wise proportion of behavioural performance explained by the interaction of predictability and intelligibility already used to relate BOLD activity to behavioural performance. To match the number of DCM parameter estimates, we averaged behavioural data according to the binning scheme described above, thus yielding six behavioural values (predictability: high, low; intelligibility: high, medium, low). Those connections with a significant intelligibility-by-predictability interaction of modulatory parameters were correlated with behavioural performance across binned conditions at the single-participant level. Fisher’s z-transformed Pearson product-moment correlation coefficients were tested against zero with a one-sample *t*-test. This analysis was run in Matlab using a custom script.

## Results

### Semantic predictability benefits speech comprehension at intermediate levels of intelligibility

On average, participants repeated 52.33% of keywords correctly (*SD* = 8.09). The threshold of psychometric curves fitted to the proportion of correctly repeated keywords across intelligibility levels decreased for sentences with high predictability when compared to low predictability (*t*_25_ = -9.87, *p* < 0.001, *r* = 0.57, BF_10_ > 1,000; Figure 2A). This threshold reflected the facilitatory effect that high semantic predictability had on speech comprehension at intermediate intelligibility levels. Moreover, the lapse rate was smaller for sentences with high compared to low predictability (*Z* = -2.78, *p* = 0.005, *r* = 0.55), indicating a predictability gain that persisted (to a smaller degree) beyond intermediate intelligibility in the highest intelligibility level.

**Figure 2.**
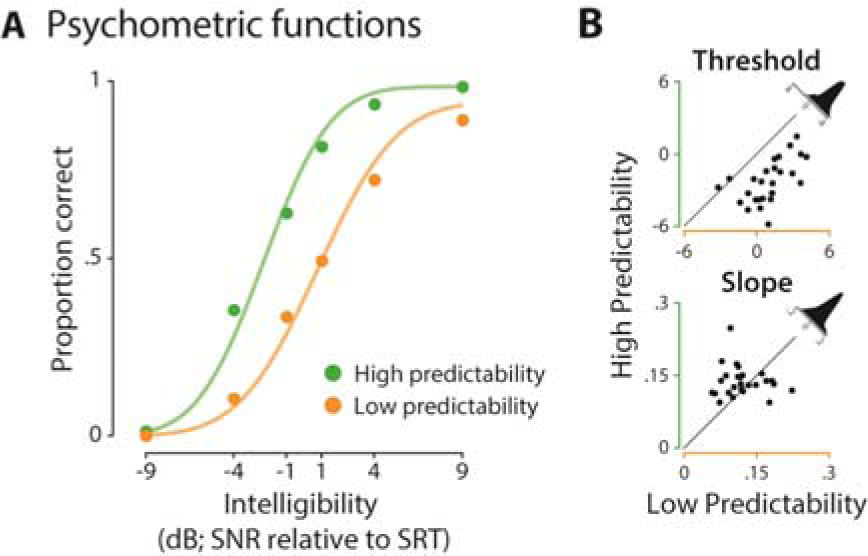
Behavioural results of the sentence repetition task. **(A)** Psychometric functions (coloured lines) were fitted to the proportion of correctly repeated keywords (coloured dots) across intelligibility levels for sentences with low (orange) and high predictability (green); grand average is displayed for illustration purposes. **(B)** The threshold of sentences with low predictability shifted towards more intelligible levels when compared to sentences with high predictability (*p* < 0.001; top). There was no evidence for a slope difference between sentences with low and high predictability (*p* = 0.122; bottom). Density plots illustrate the difference between the proportions correct of sentences with high vs. low predictability.

There was no significant difference between sentences with low and high predictability for slope (*t*_25_ = 1.6, *p* = 0.122, *r* = -0.11, BF_10_ = 0.64), width (*t*_25_ = 1.84, *p* = 0.077, *r* = 0.11, BF_10_ = 0.896) and guess rate (*Z* = 0.27, *p* = 0.79, *r* = 0.05) of psychometric curves.

Importantly, even though SRTs from the adaptive tracking procedure were strongly correlated with the overall proportion of correctly repeated keywords (*r* = 0.89, *p* < 0.001, BF_10_ > 1,000; see Supplementary Figure 6), effects of predictability on psychometric curves were unaffected by these inter-individual differences in auditory perception. When controlling for the potentially confounding influence of individual SRTs on parameter estimates, effects of threshold (*t*_25_ = -9.31, *p* < 0.001, *r* = 0.13, BF_10_ > 1,000) and lapse rate (*t*_25_ = -2.09, *p* = 0.047, *r* = 0.06, BF_10_ = 1.33) remained significant and all other effects remained non-significant.

### A fronto-temporo-parietal network is tuned towards intelligible speech

First, we were interested in brain regions showing stronger activity for increasing intelligibility of speech stimuli, irrespective of sentence predictability. The parametric variation of intelligibility showed increased activation in a set of regions comprising anterior-to-posterior bilateral superior temporal gyri (STG), left precentral and postcentral gyrus and left inferior frontal gyrus (IFG) as well as a large left-hemispheric temporo-parietal cluster and left precuneus (Figure 3). These regions have previously been reported to support auditory speech comprehension and suggested to be tuned to perceptual clarity (Rauschecker & Scott, 2009).

**Figure 3.**
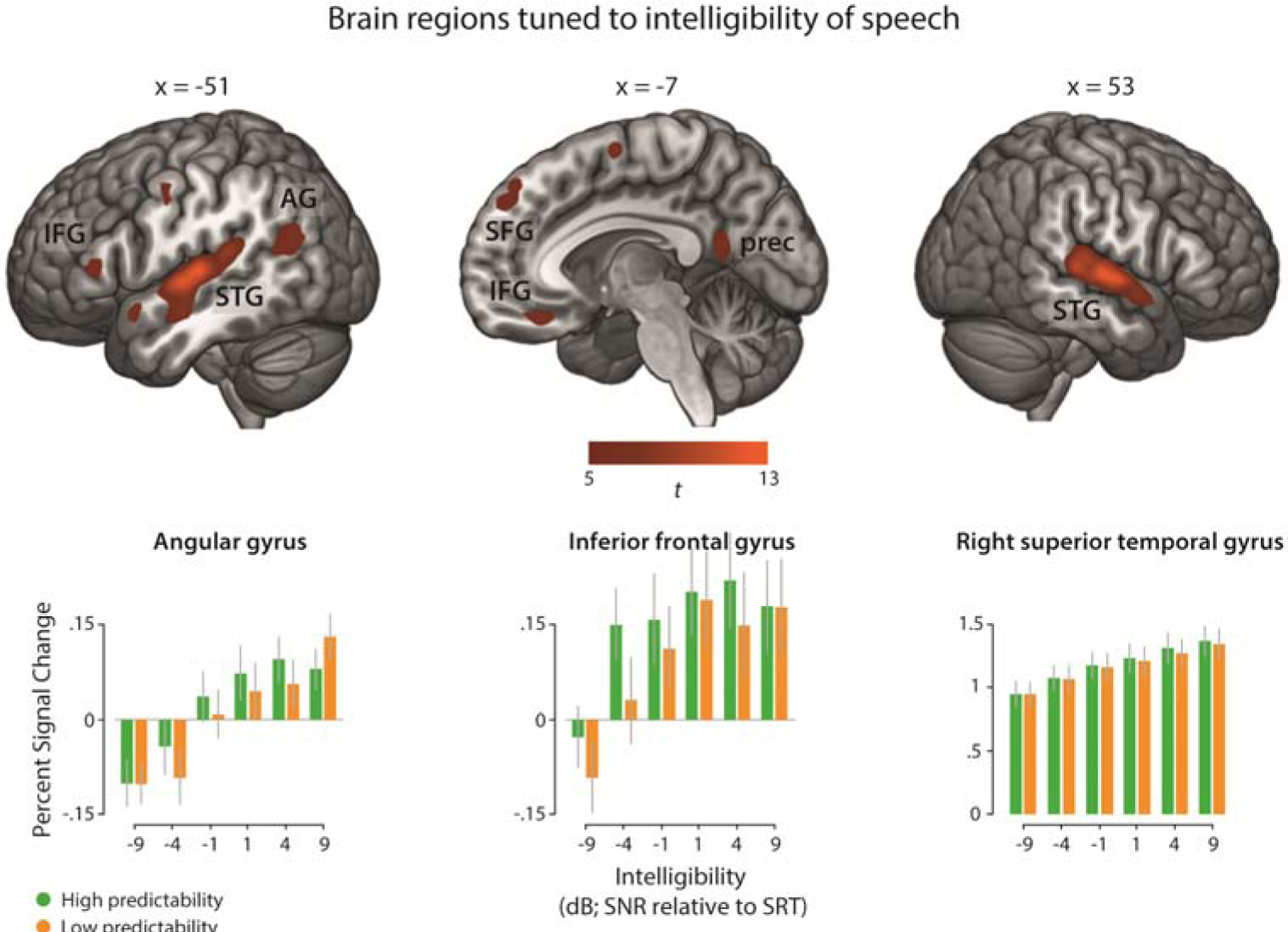
Main effect of increasing speech intelligibility across the whole cortex. Top: Activation map thresholded at *p* < 0.05 (FWE-corrected). STG = superior temporal gyrus, IFG = inferior frontal gyrus, AG = angular gyrus, SFG = superior frontal gyrus, prec = precuneus. Bottom: Average parameter estimates (percent signal change) for each experimental condition at peak voxels of left AG (MNI: x = -47, y = -67, z = 25), left IFG (MNI: x = -54, y = 33, z = 8) and right STG (MNI: x = 53, y = -12, z = 2). Error bars represent the standard error of the mean (SEM).

### Increasing intelligibility of high predictable sentences enhances recruitment of left-hemispheric temporo-parietal regions

Further, we were interested in brain regions that were differentially affected by predictability at different levels of intelligibility. Higher activation for sentences with high compared to low predictability under linearly increasing intelligibility was found in left-hemispheric regions encompassing pMTG, middle to posterior cingulate cortex, two separate clusters in left AG (PGa and PGp), inferior temporal gyrus, precuneus and supramarginal gyrus (SMG; Figure 4, Table 1). Similar but weaker activation patterns were found for the quadratically modulated interaction contrast. Interestingly, the two separate left-hemispheric AG clusters fell into different cytoarchitectonic subregions: the larger cluster was mainly located in the anterior dorsal portion of AG (PGa & PFm) at the boundary to SMG and the smaller cluster was mainly located in the posterior portion of AG (PGp) in close proximity to the middle occipital gyrus (MOG). The AG clusters and the pMTG cluster showed a pattern of deactivation relative to the fixation baseline with relatively less deactivation for high predictable as compared to low predictable sentences (Figure 4B). As both, the AG and pMTG are considered key regions of semantic processing, their time series were submitted to later connectivity analyses.

**Figure 4.**
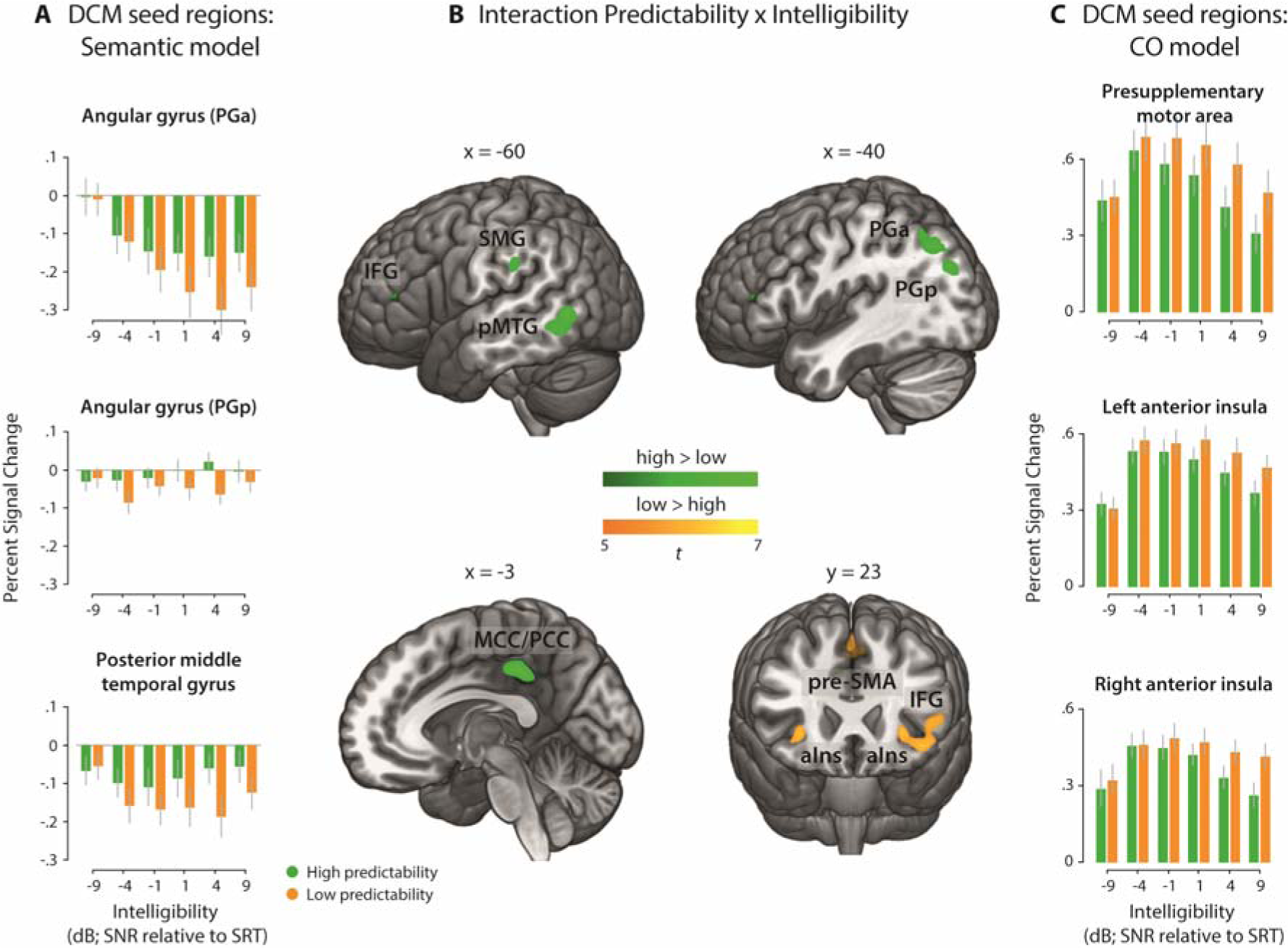
Group-level fMRI results illustrating brain regions that are sensitive to the interaction of intelligibility and predictability. **(A)** Parameter estimates illustrate the average interaction effect, extracted from peak voxels of DCM seed regions for the semantic network. **(B)** Brain regions showing a significant interaction; IFG = inferior frontal gyrus, SMG = supramarginal gyrus, pMTG = posterior middle temporal gyrus, PGa = anterior angular gyrus, PGp = posterior angular gyrus, MCC/PCC = middle and posterior cingulate cortex, pre-SMA = pre-supplementary motor area, aIns = anterior insula; green = high > low predictable sentences, orange = low > high predictable sentences. The activation map is thresholded at *p*_FWE-corrected_ < 0.05. **(C)** Parameter estimates illustrate the interaction effect, extracted from peak voxels of DCM seed regions for the cingulo-opercular (CO) network. Error bars represent ±1 SEM.

**Table 1.** Results from the context-dependent interaction.

| Location | Hemisphere | MNI |  |  | t | Cluster Size |
| --- | --- | --- | --- | --- | --- | --- |
|  |  | x | y | z |  |  |
| High > Low predictable sentences |  |  |  |  |  |  |
| Middle cingulate cortex | R | 3 | −34 | 38 | 7.49 | 159 |
| Middle cingulate cortex | L | −4 | −30 | 42 | 7.17 | subcluster |
| Middle temporal gyrus | L | −60 | −52 | −5 | 6.85 | 150 |
| Middle temporal gyrus | L | −54 | −62 | 2 | 6.6 | subcluster |
| Angular gyrus (PFm/PGa) | L | −42 | −62 | 50 | 6.47 | 135 |
| Angular gyrus (PGa) | L | −42 | −70 | 42 | 6.31 | subcluster |
| Angular gyrus (PGp) | L | −34 | −77 | 42 | 6.02 | subcluster |
| Angular Gyrus (PGp)/<br>middle occipital gyrus | L | −34 | −77 | 30 | 6.34 | 51 |
| Angular gyrus (PGp) | L | −42 | −77 | 28 | 5.63 | subcluster |
| Supramarginal gyrus | L | −60 | −30 | 30 | 5.91 | 26 |
| Inferior temporal gyrus | L | −47 | −52 | −25 | 5.82 | 21 |
| Precuneus | L | −10 | −47 | 55 | 5.62 | 15 |
| Low > High predictable sentences |  |  |  |  |  |  |
| Presupplementary motor<br>area/paracingulate gyrus | R/L | 0 | 16 | 50 | 7.72 | 138 |
| Inferior frontal gyrus (p. orbitalis,<br>area 45) | L | −42 | 26 | −10 | 6.47 | 106 |
| Insular cortex | L | −30 | 26 | −2 | 6.43 | subcluster |
| Inferior frontal gyrus (p. triangularis,<br>area 45) | L | −50 | 23 | 5 | 5.9 | subcluster |
| Inferior frontal gyrus (p. triangularis,<br>area 45) | L | −44 | 20 | 0 | 5.88 | subcluster |
| Insular cortex | R | 30 | 26 | −2 | 6.13 | 24 |
| Inferior frontal gyrus (p. triangularis,<br>BA 45) | L | −57 | 20 | 15 | 5.71 | 16 |
| Inferior frontal gyrus (p. triangularis,<br>BA 45) | L | −50 | 20 | 18 | 5.07 | subcluster |

High > low predictable sentences: Regions showing greater activity for high as compared to low predictable sentences under increasing intelligibility; Low > high predictable sentences: Regions showing greater activity for low as compared to high predictable sentences under increasing intelligibility. Regions are reported at a peak-level FWE-correcting threshold at an alpha of .05. Seed regions for DCM analyses are highlighted in bold.

### Increasing intelligibility of low predictable sentences enhances recruitment of domain-general regions

The opposite interaction contrast revealed greater activity for low compared to high predictable sentences under linearly increasing intelligibility and encompassed pre-SMA/paracingulate gyrus, left-hemispheric IFG (pars triangularis; BA 45) and the bilateral anterior insula (see Figure 4). These regions overlap with the cingulo-opercular network (Dosenbach et al., 2008), which is frequently reported in conditions associated with high task demands. To investigate context-dependent changes within this network, the connectivity between pre-SMA/paracingulate gyrus and bilateral insulae was analysed in a later DCM analysis. Note that the interaction contrasts modelling intelligibility quadratically (i.e. inverted u-shape with the highest activation at intermediate levels of intelligibility) resulted in a similar pattern of brain regions and therefore are not shown here.

### The extended semantic system scales to the behavioural predictability gain dependent on speech intelligibility

The proportion of individual task performance explained by the interaction of predictability and intelligibility in each experimental condition correlated positively with BOLD activity in a broad semantic network (Figure 5; Binder et al., 2009). At intermediate levels of intelligibility, neural activity increased for sentences with high predictability in those participants exploiting a stronger behavioural predictability gain.

**Figure 5.**
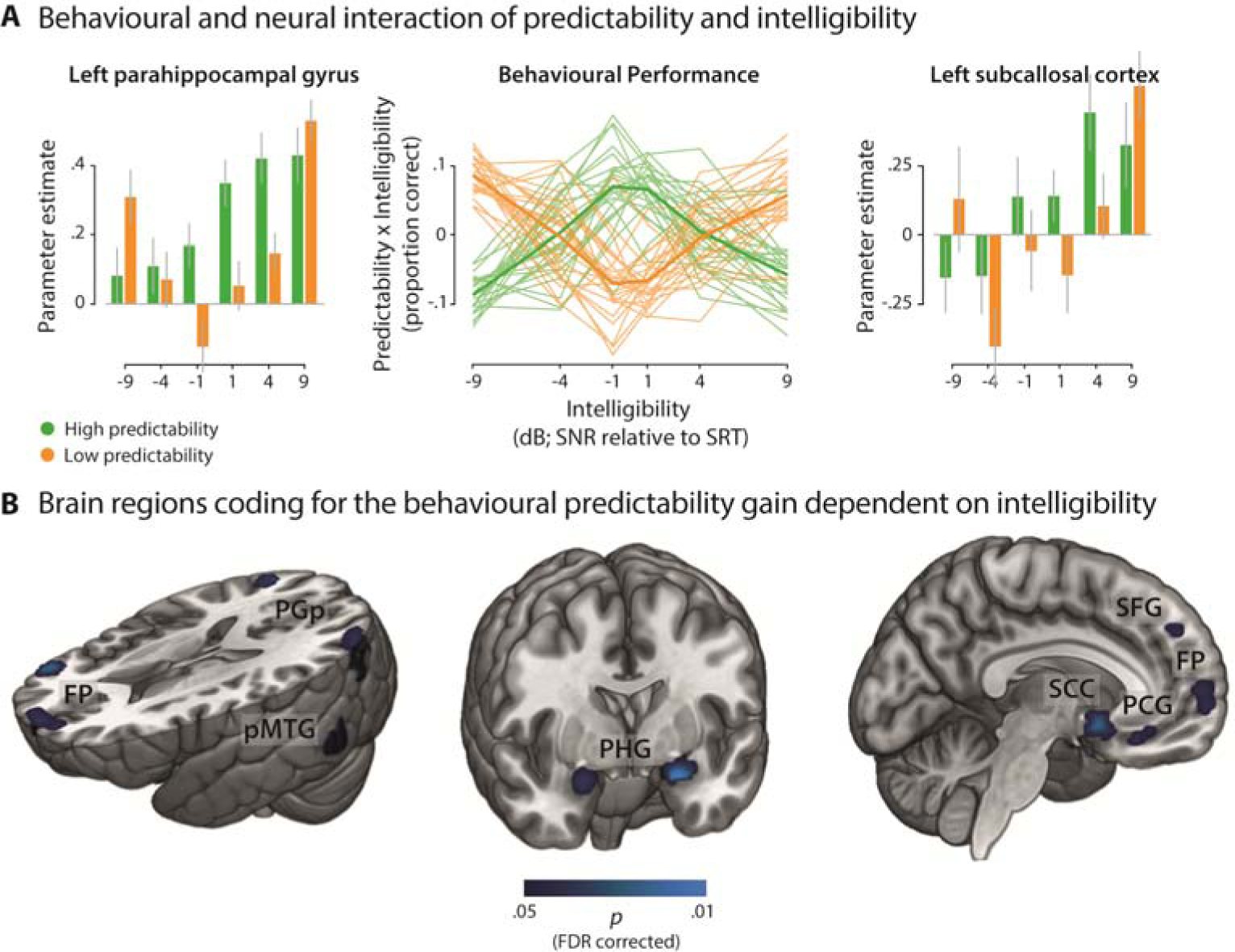
Projection of activity onto the behavioural predictability gain across levels of intelligibility. **(A)** The condition-specific proportion of behavioural performance explained by the interaction of intelligibility and predictability in each participant (middle line plot) was correlated with individual parameter estimates of single-voxel BOLD activity (left and right bar plots; exemplary grand average parameter estimates for two peak voxels). Thin lines in the line plot represent single participants; fat lines represent grand average. Error bars represent ±1 SEM. **(B)** Group statistics revealed brain areas modulated by the single-participant interaction effect of intelligibility and predictability on speech comprehension. *P*-values were FDR-corrected within a mask of voxels responsive to the listening task. FP = frontal pole; PGp = posterior angular gyrus; pMTG = posterior middle temporal gyrus; PHG = parahippocampal gyrus; SCC = subcallosal cortex; SFG = superior frontal gyrus; PCG = paracingulate gyrus (for an exhaustive list of clusters see Table 2).

**Table 2.**
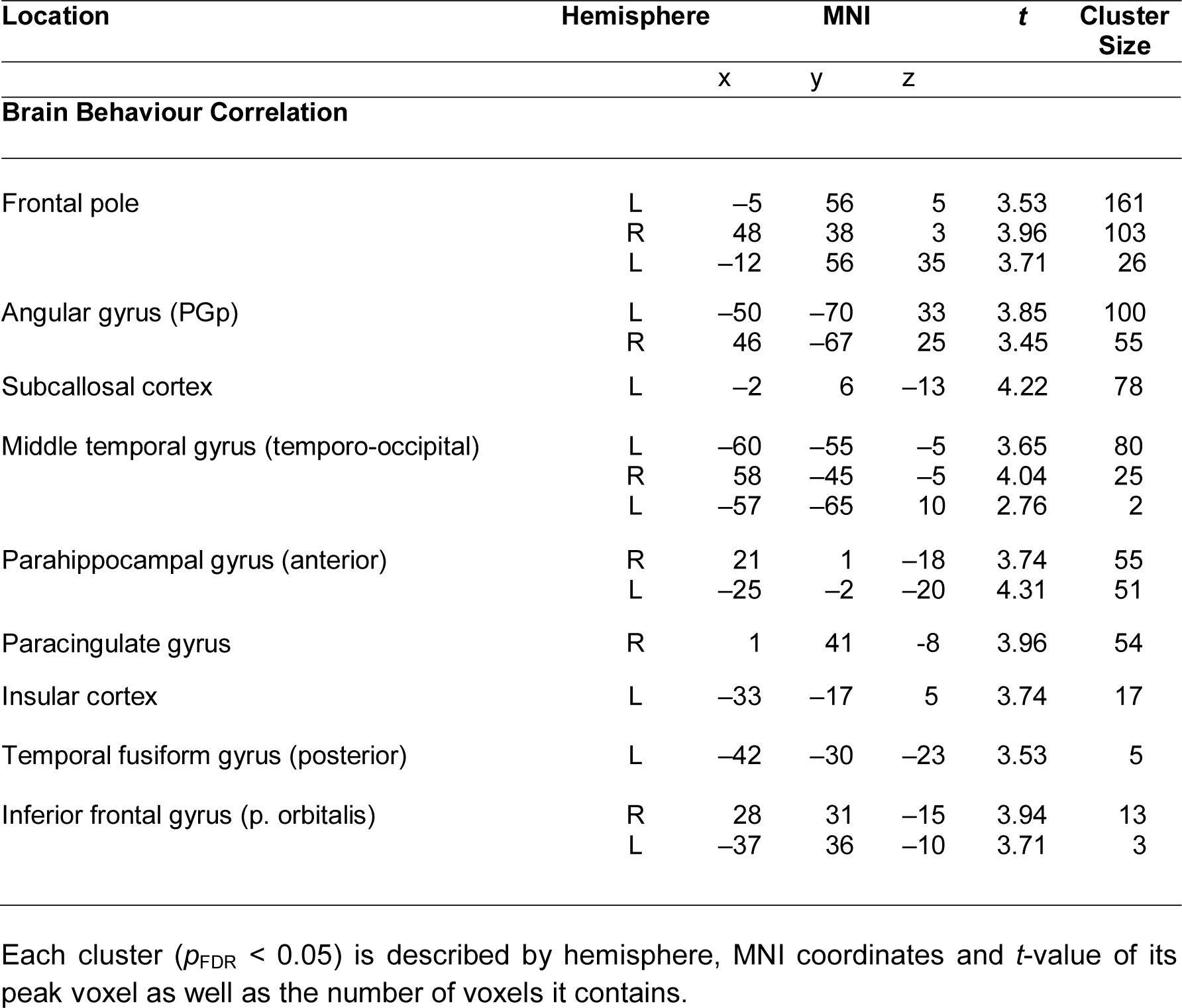
Brain areas sensitive to the interaction effect of intelligibility and predictability on task performance.

| Location | Hemisphere | MNI |  |  | t | Cluster Size |
| --- | --- | --- | --- | --- | --- | --- |
|  |  | x | y | z |  |  |
| Brain Behaviour Correlation |  |  |  |  |  |  |
| Frontal pole | L | −5 | 56 | 5 | 3.53 | 161 |
|  | R | 48 | 38 | 3 | 3.96 | 103 |
|  | L | −12 | 56 | 35 | 3.71 | 26 |
| Angular gyrus (PGp) | L | −50 | −70 | 33 | 3.85 | 100 |
|  | R | 46 | −67 | 25 | 3.45 | 55 |
| Subcallosal cortex | L | −2 | 6 | −13 | 4.22 | 78 |
| Middle temporal gyrus (temporo-occipital) | L | −60 | −55 | −5 | 3.65 | 80 |
|  | R | 58 | −45 | −5 | 4.04 | 25 |
|  | L | −57 | −65 | 10 | 2.76 | 2 |
| Parahippocampal gyrus (anterior) | R | 21 | 1 | −18 | 3.74 | 55 |
|  | L | −25 | −2 | −20 | 4.31 | 51 |
| Paracingulate gyrus | R | 1 | 41 | −8 | 3.96 | 54 |
| Insular cortex | L | −33 | −17 | 5 | 3.74 | 17 |
| Temporal fusiform gyrus (posterior) | L | −42 | −30 | −23 | 3.53 | 5 |
| Inferior frontal gyrus (p. orbitalis) | R | 28 | 31 | −15 | 3.94 | 13 |
|  | L | −37 | 36 | −10 | 3.71 | 3 |

Each cluster ( $p_{\text{FDR}} < 0.05$ ) is described by hemisphere, MNI coordinates and *t*-value of its peak voxel as well as the number of voxels it contains.

In line with the linear interaction effect (see Table 1, high > low predictability), modelling the behavioural interaction effect revealed significant clusters in parietal and lateral temporal cortex. A broad left-hemispheric and a smaller right-hemispheric cluster were observed in PGp (Table 2). Additional clusters were found in bilateral pMTG.

More importantly, the behavioural interaction effect mapped onto frontal and ventromedial brain regions not implicated in the linear interaction effect. The frontal pole encompassed one broad medial cluster spanning from the left to the right hemisphere as well as a right-hemispheric cluster at the border with IFG and a left-hemispheric cluster extending into the superior frontal gyrus (Table 2). Near the inferior-posterior frontal pole, we observed bilateral clusters in IFG. The limbic lobes contained a bilateral cluster in the posterior subcallosal cortex that was complemented by a bilateral cluster in inferior paracingulate gyrus extending into anterior subcallosal cortex as well as cingulate gyrus and frontal medial cortex. Additional clusters comprised the anterior division of bilateral superior parahippocampal gyrus and left medial-posterior insular cortex. In the medial temporal lobe, we found a left-hemispheric cluster in the posterior division of the temporal fusiform gyrus. No significant cluster in the brain showed a negative correlation with the interaction effect on task performance.

### Connectivity between subregions of the angular gyrus increases with higher intelligibility

Next, to investigate context-dependent changes in effective connectivity between regions identified in the mass univariate GLM analysis, we performed two separate DCM analyses, one within the semantic system and one within the cingulo-opercular network.

The family of models with a modulatory influence on the connection from PGp to pMTG and PGa was identified as the winning family by means of Bayesian model selection (exceedance probability: 0.65; see Supplementary Figure 7A for an overview of all exceedance probabilities). All intrinsic connections, except for self-connections, were positive (see Supplementary Table 1), although only the outgoing connections from pMTG and the self-connections reached statistical significance. Within the winning family, there was a significant main effect of intelligibility on the connection from PGp to PGa (*F*_1.56, 39.07_ = 3.57, *p* = 0.048, 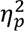 = 0.13, Greenhouse-Geisser corrected, Mauchly’s sphericity test: *p* = 0.02). The modulatory influence of intelligibility increased connectivity from PGp to PGa when speech became more intelligible. The main effect of predictability (*F*_1, 25_ = 1.86, *p* = .185, 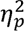 = .07) and the interaction effect (*F*_2, 50_ = 0.71, *p* = .496, 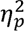 = 0.03) were not significant. There was no significant effect on the connection from PGp to pMTG (main effect predictability: *F*_1, 25_ = 0.02, *p* = .899, 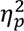= .001; main effect intelligibility: *F*_2, 50_ = 0.35, *p* = .705, 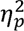 = .01; interaction effect: *F*_2, 50_ = 1.55, *p* = .222, 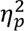 = 0.06).

### Inhibitory connectivity within the cingulo-opercular network increases with the individual predictability gain at intermediate intelligibility

For the cingulo-opercular network model, the family of models with a modulatory effect on the connections from pre-SMA and right insula to left insula was identified as the winning family, with an exceedance probability of 0.73 (see Supplementary Figure 7B).

In the winning model, there was a significant interaction in the connection from pre-SMA to left insula (F_2,50_ = 4.74, p = 0.013, 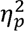 = 0.016): inhibitory influence was less pronounced for low predictable sentences with increasing intelligibility but stronger for high predictable sentences with increasing intelligibility. The respective main effects of intelligibility (*F*_2, 50_ = 0.55, *p* = .583, 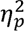 = .02) and predictability (*F*_1, 25_ = 0.11, *p* = .747, 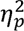 = .004) were not significant (see Figure 6A and Supplementary Table 2). There was no significant effect on the connection from right insula to left insula (main effect intelligibility: *F*_2, 50_ = 0.36, *p* = .702, 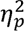 = .01; main effect predictability: *F*_1, 25_ = 1.64, *p* = .212, 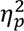 = .06; interaction effect: *F*_2, 50_ = 2.07, *p* = .137, 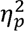 = 0.08).

**Figure 6.**
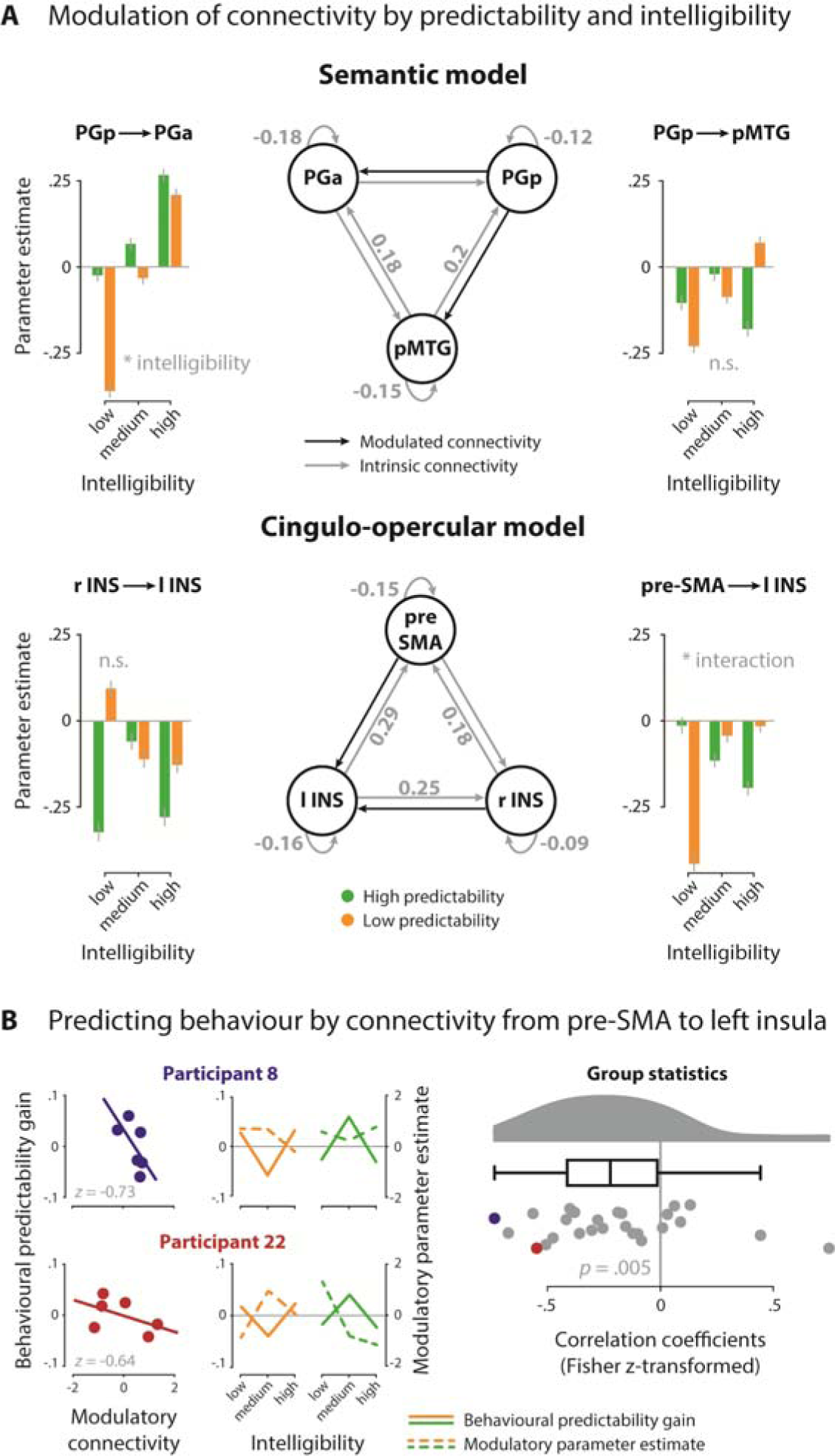
DCM results. **(A)** Network architecture of the semantic (top) and cingulo-opercular network (bottom); grey arrows with parameter estimates indicate significant intrinsic connections; black arrows indicate modulatory connections. Bar graphs show the modulation of connections by intelligibility (low, medium, high; binned) and predictability (low, high). The connection from PGp to PGa was significantly modulated by the main effect of intelligibility, the connection from pre-SMA to left insula was sensitive to the interaction of predictability and intelligibility. Bar height represents grand average, error bars represent ±1 SEM. **(B)** Scatterplots (left) show the negative correlation of the condition-wise predictability gain with the condition-wise modulatory parameter estimates from pre-SMA to left insula in two exemplary participants. Complementary, line plots (middle) illustrate patterns of behavioural performance (solid line) and modulation strength (dashed line) resolved for intelligibility (low, medium, high) and predictability (low = orange; high = green). Raincloud plot (right) shows the distribution of individual *z*-transformed correlations across all participants; dots representing participants from scatter and line plots are highlighted in colour. n.s. = not significant, * *p* < 0.05. PGa = anterior angular gyrus; PGp = posterior angular gyrus; pMTG = posterior middle temporal gyrus; pre-SMA = pre-supplementary motor area; l INS = left anterior insula; r INS = right anterior insula.

Following up the interaction effect in connectivity between pre-SMA and left insula with a post-hoc analysis, we investigated how individual connectivity patterns match the individual pattern of the behavioural predictability gain across intelligibility levels. We found a significant negative correlation (*t*_25_ = -3.919, *p* = 0.004, BF_10_ = 53.17), indicating that the individual predictability gain at medium intelligibility was associated with stronger inhibitory influence from pre-SMA to the left insula for highly predictable speech (see Figure 6B).

## Discussion

In the present fMRI study, we investigated the neural underpinnings of the semantic predictability gain during speech processing in noisy listening situations at the network level. First, we showed that highly predictable but acoustically degraded speech engages left-hemispheric semantic regions and modulates the individual comprehension gain in behaviour via additional frontal and medial semantic regions. Second, we found that highly intelligible speech strengthens effective connectivity between two subregions of a key semantic network node in the left angular gyrus. Third, we showed for the cingulo-opercular network that inhibitory influence from pre-SMA to left insula was stronger for increasing intelligibility of highly predictable speech but was less pronounced for low predictability. Notably, individual behavioural response patterns were negatively correlated with individual connectivity patterns. Specifically, the higher the behavioural comprehension gain at intermediate intelligibility was, the stronger the inhibitory connectivity strength within individual subjects. These findings suggest that semantic predictability facilitates speech comprehension in challenging listening conditions via upregulation of the semantic network and active inhibition between domain-general regions.

Our sentence repetition task was designed to overcome the experimental shortcomings of previous studies and investigate speech comprehension in noise on the basis of rich neural and behavioural data. Hitherto, fMRI studies on the neural underpinnings of degraded speech processing had mainly used sparse temporal sampling to avoid interference of scanner noise with experimentally manipulated signal-to-noise ratio of speech stimuli (Erb et al., 2013; Golestani et al., 2013; Obleser & Kotz, 2010). As we aimed to investigate speech processing directly during (compared to after) listening and to improve sampling density for our connectivity analyses, BOLD activity in the present study was continuously recorded throughout the whole experiment. In line with studies using temporal sparse sampling (Abrams et al., 2013; Davis & Johnsrude, 2003; Rauschecker & Scott, 2009), we found activity in regions associated with auditory comprehensibility (e.g., bilateral STG as well as left IFG) to be stronger with increasing intelligibility of speech. This replication of the intelligibility effect speaks for continuous scanning as a feasible technique to investigate *online* speech processing.

Crucially, in contrast to previous studies with only few and fixed experimental levels of speech degradation, we covered the full range of intelligibility from hardly to easily comprehensible speech in the individual participant. Note that our experimental design contained neither a “noise only” nor a clear speech condition. At the behavioural level, we were nevertheless able to fit full psychometric curves (i.e., speech comprehension as a function of intelligibility, see Figure 2). We found the psychometric curve to shift towards less intelligible levels for sentences with high compared to low semantic predictability. This threshold shift is an established effect in more challenging comprehension tasks (Shannon et al., 2004) and reflects a comprehension benefit for predictable speech at intermediate intelligibility.

As left AG has been found to show a strong differentiation between sentences with high and low predictability at intermediate intelligibility (Obleser et al., 2007; Obleser & Kotz, 2010) and to exhibit functional relevance in mediating the comprehension gain (Hartwigsen et al., 2015), this region has been proposed to serve top-down activation of semantic concepts that facilitate speech comprehension (Obleser & Kotz, 2010). However, as an unexpected finding, two separate AG clusters showed stronger activation for sentences with high compared to low predictability under linearly increasing intelligibility: a larger cluster mainly falling into PGa at the boundary to the supramarginal gyrus, and a smaller cluster at the boundary between PGp and middle occipital gyrus. Importantly, these AG subregions have been discussed to subserve different functions (Bonnici et al., 2016; Noonan et al., 2013; Seghier et al., 2010). PGa is implicated in domain-general attention and converges with coordinates commonly reported as AG activation during speech in noise processing (Clos et al., 2014; Guediche et al., 2014; Obleser et al., 2007; Obleser & Kotz, 2010). In contrast, PGp activation has often been labelled as MOG (Guediche et al., 2016; McGettigan et al., 2012). In these studies, MOG was interpreted to provide free resources otherwise recruited by visual processing to the auditory domain during resource-demanding speech in noise comprehension. Other studies have associated PGp activation with “pure” semantic processing (Bonnici et al., 2016; Seghier et al., 2010). Our findings implicate that both AG subregions contribute to successful speech comprehension under challenging listening conditions when semantic cues are available.

To further characterize the functional contributions of anterior and posterior AG, we analysed the connectivity between these subregions and observed that coupling from posterior to anterior AG was strengthened when speech became more intelligible. Surprisingly, this connection was not modulated by the predictability of speech. This finding suggests that the interplay of AG subregions might support the integration of lexical information in general, but does not specifically facilitate integration of semantic information. Further, this finding speaks against the view that the anterior portion (PGa) is associated with goal-directed attention processes as one would expect attentional demands to be highest at intermediate levels of intelligibility and not at the most intelligible levels. However, it is important to bear in mind that the effective connectivity results are restricted to the limited set of regions included in the network models. Aside from AG, we largely found the same regions (e.g., left pMTG, left SMG, left inferior temporal gyrus, posterior cingulate gyrus and precuneus) that were previously implicated in processing of sentences with high predictability under increasing intelligibility (Golestani et al., 2013; Obleser et al., 2007; Obleser & Kotz, 2010). These regions overlap with the previously described core semantic (control) regions (Binder et al., 2009; Jefferies, 2013) and form a functional network during degraded speech processing (Obleser et al., 2007).

To complement the linear interaction contrast, we also modelled task performance in the brain and expected to find the left-hemispheric semantic network strengthened by this contrast fine-tuned to inter-subject variation. Indeed, the semantic core regions extended to their homologous regions in the right hemisphere. Strikingly, additional semantic regions scaled in activation with the behavioural predictability gain at intermediate intelligibility. These regions largely pertained to two subsets of the semantic network: *lateral and ventral temporal cortex* includes left temporal fusiform gyrus and parahippocampal gyrus, whereas *left ventromedial prefrontal cortex* includes subcallosal cortex and frontal pole (Binder et al., 2009). The temporal cortex has been shown to communicate with the hippocampus via a connection from temporal fusiform gyrus to parahippocampal gyrus evident in structural (Powell et al., 2004) and functional imaging (Teipel et al., 2010). It has been suggested that this pathway is used to encode semantic information (Levy et al., 2004; Martin, 2007). Critically, ventral temporal activation has been implicated in the processing of semantically unambiguous speech in a memory-demanding word recognition task (Rodd et al., 2005) and processing of visually presented words that were correctly repeated during later recall (Strange et al., 2002). In the sentence repetition task employed here, memory formation is crucial to correctly repeat the sentence after listening. The association of more accurate recall with stronger ventral temporal activation for predictable but degraded sentences suggests that memory engagement is most crucial for successful speech comprehension when semantic information is available for integration and must be protected against competing but irrelevant information (i.e., background noise). In line with this notion, prefrontal cortex regions implicated in executive control functioning (Duncan & Owen, 2000) show increased activation when efficient management of cognitive resources promises the strongest gain in performance. While we will refrain from overinterpreting these exploratory results, it is worth highlighting that the role of executive control and memory formation might have been often overlooked in previous studies on speech comprehension despite their face validity when it comes to understanding individual hearing difficulties.

As a last key finding of our study, challenging speech comprehension in the absence of semantic constraints is accompanied by an increase of activity in the pre-SMA and bilateral anterior insulae, which are regions associated with the cingulo-opercular network. This finding is broadly in line with previous reports on activation of cingulo-opercular regions during effortful speech processing (Adank, 2012; Clos et al., 2014; Eckert et al., 2009; Erb & Obleser, 2013; Hervais-Adelman et al., 2012; Vaden et al., 2013). The cingulo-opercular network has been associated with domain-general control or executive processes such as task monitoring (e.g. Dosenbach et al., 2008; Duncan, 2010; Vaden et al., 2013, 2015),executive control (Erb & Obleser, 2013) and attentional control (Fitzhugh et al., 2019). Specifically, in the speech in noise literature, the increased reliance on this network in difficult listening conditions has been interpreted as an adaptive control mechanism to enable speech comprehension when little or no semantic information is available.

Within these cingulo-opercular regions, we found an inverted u-shaped response, with highest activation at intermediate levels of intelligibility and less activation for the conditions that were either hardly or easily comprehensible. This activation pattern suggests that domain-general regions contribute to effortful speech processing as long as *any* information can be extracted. Analogously, recruitment of these regions is less important when the auditory signal is too bad or when speech is clear. On the other hand, the insula has been shown to mirror the inverse quadratic response time effect across intelligibility levels, thereby reflecting the involvement of the insula in response selection (Binder et al., 2004). Even though we did not use a discrimination task but a sentence repetition paradigm, the number of lexical candidates for the response might be highest when speech is hardly comprehensible and not guided by semantic information, thereby explaining an increase in insula activity. However, this does not explain the consistent response pattern across different regions implicated in the cingulo-opercular network that are typically not involved in response selection. Moreover, in a recent meta-analysis, bilateral insulae were linked to higher-order cognitive aspects of speech comprehension and production, highlighting that insula function goes beyond mere involvement in motor aspects of speech production (Oh et al., 2014).

Further, we observed that connectivity between these regions was differentially modulated by high and low predictability. These findings suggest that active inhibition is increased between domain-general cognitive control regions when predictability and intelligibility were highest or lowest. Interestingly, effective connectivity from pre-SMA to the left insula was associated with individual task performance, that is, the inhibitory modulation of this connection increased with the individual degree of the predictability gain at intermediate intelligibility (and decreased with the relative behavioural loss from low predictability). This result converges with and extends previous task-based fMRI studies that found increased activity in the bilateral insulae when auditory input was degraded (Erb et al., 2013; Hervais-Adelman et al., 2012). While older adults with well-preserved hearing abilities and younger participants recruited the anterior insula under challenging listening conditions, recruitment of this region was already observed in clear speech conditions for older adults with hearing loss (Erb & Obleser, 2013). Together, the previous and present findings suggest that recruitment of the left anterior insula is beneficial in challenging conditions (e.g. when auditory input is deteriorated) but becomes unnecessary as soon as the task is easily or not at all solvable. Moreover, the present findings suggest that better behavioural performance may require an active inhibition of this region to ensure efficient processing.

Recently, the pre-SMA has been suggested to be relevant during high task demands (Hertrich et al., 2016) and to play a role in the wider semantic control network (Hallam et al., 2016). Specifically, Hallam and colleagues (2016) found increased activity in pre-SMA and pMTG after inhibitory TMS to left IFG, which was interpreted as upregulation of the relative contribution to semantic control in the presence of a TMS-induced disruption. Further, Dietrich and colleagues (2018) reported decreased performance in a sentence repetition task after inhibitory TMS over the pre-SMA, which was selectively evident for the most challenging task condition. Note that pre-SMA activation has also been reported in sentence comprehension tasks in the degraded speech literature before (e.g. Clos et al., 2014: sentence matching task). Therefore, it is unlikely that our pre-SMA finding mainly reflects the choice of the task alone. Taken together, our results replicate the previously reported involvement of domain-general regions during effortful speech processing and extend these findings by demonstrating predictability-dependent modulation of functional connectivity within this network.

## Conclusion

Our results demonstrate that predictability in challenging listening conditions not only modulates the semantic network but critically extends to modulations in the domain-general network in a behaviourally relevant fashion. We demonstrated that intelligibility modulated connectivity between two subregions of left AG, underscoring the functional heterogeneity of this region. Further, the degree of inhibitory modulation of connectivity within the domain-general cingulo-opercular network was associated with the individual speech comprehension gain in behaviour. This highlights the importance to further investigate the dynamic interplay between the semantic and cingulo-opercular network in successful speech comprehension under adverse listening conditions.

## Acknowledgements

The authors would like to thank Manuela Hofmann, Sylvie Neubert and Simone Wipper for support with the data acquisition and Dagny Kühner, Isabel Gebhardt, Anna Ruhe, and Ricarda Vielhauer for support with recording transcription. We further thank Julia Erb for providing the stimulus materials and the University of Minnesota Center for Magnetic Resonance Imaging for providing the fMRI multiband sequence software.

## Data availability

All data are available from the corresponding authors upon reasonable request.

## Funding

This work was supported by the German Research Foundation (DFG; HA 6314/4-1 and OB 352/2-1). JO is supported by an ERC Consolidator Grant (ERC-CoG-2014; grant number 646696).

## Conflict of interest

The authors declare no conflicts of interest.

## Ethics approval

All experimental procedures were approved by the local ethics committee at the Medical Faculty of the University of Leipzig (reference number 281/16-ek).

**Patient consent:** All participants gave written informed consent.

**Permission to reproduce material from other sources:** Not applicable.

**Clinical trial registration:** Not applicable.

## Notes

### Competing Interest Statement

The authors have declared no competing interest.

