## Supplementary Material for "Neural modelling of the semantic predictability gain under challenging listening conditions"

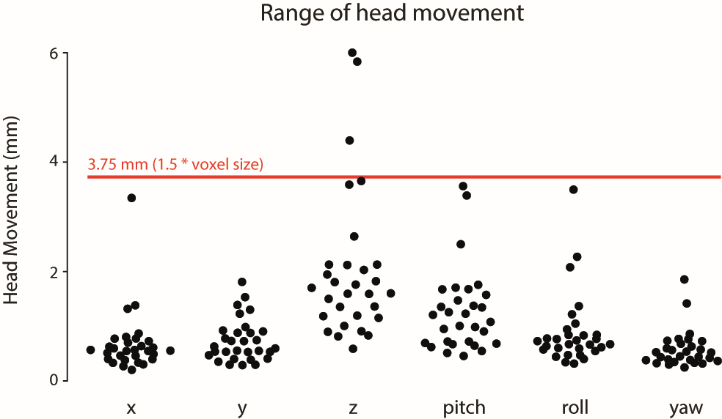

**Supplementary Figure 1.** Overview of the average head movement for each participant during the fMRI experiment. The six movement parameters were obtained from the rigid-body transformation of the realignment procedure during preprocessing and averaged for each participant. Black dots indicate individual averages for each movement parameter and the red line indicates the movement threshold that was set to 1.5 x voxel size. Participants who exceeded this threshold were excluded from further analyses.

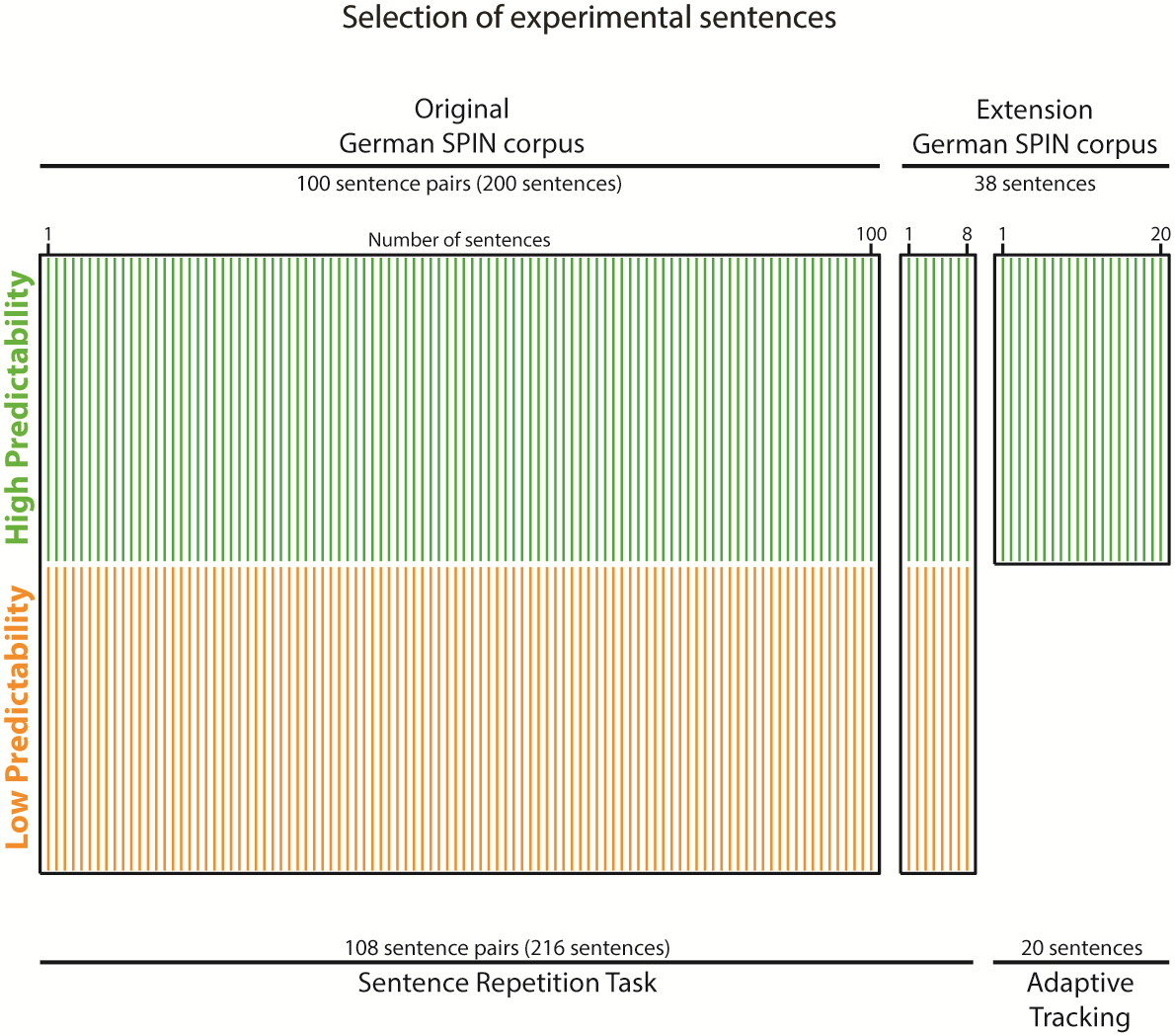

**Supplementary Figure 2.** Schematic illustration of the stimulus material. The original German SPIN corpus (Erb, Henry, Eisner, & Obleser, 2012) comprises 200 sentences with pairs of high and low predictable keywords. Together with 8 new sentence pairs, we presented 216 sentences in the sentence repetition task. We used another 20 newly rated sentences with highly predictable keywords for the adaptive tracking procedure prior to the experiment.

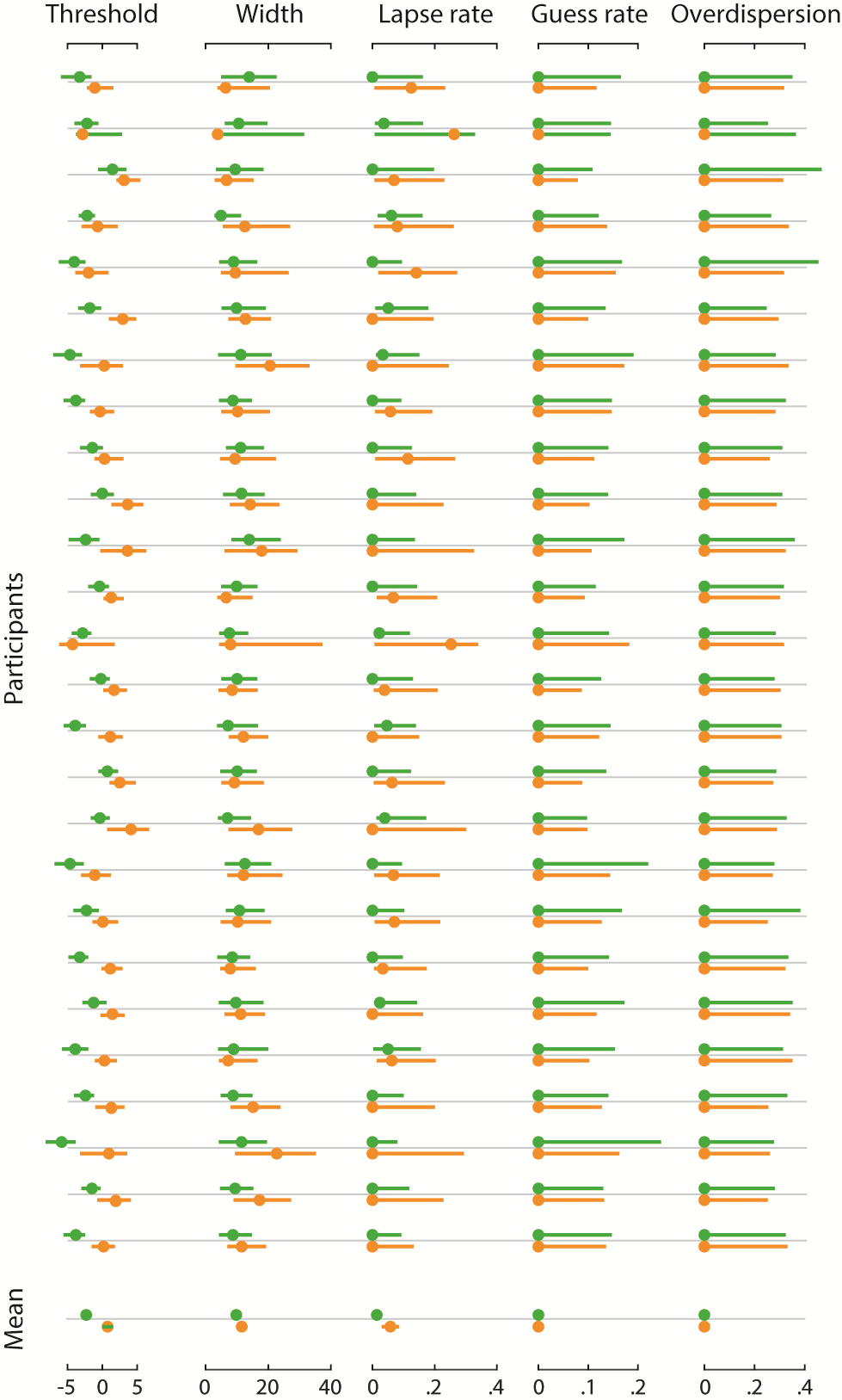

**Supplementary Figure 3.** Rows represent single-participant parameter estimates (dots) and 95 % Bayesian credible intervals (horizontal lines) of psychometric curves for sentences with low (orange) and high predictability (green). The bottom row shows parameter estimates averaged across participants with frequentist confidence intervals.

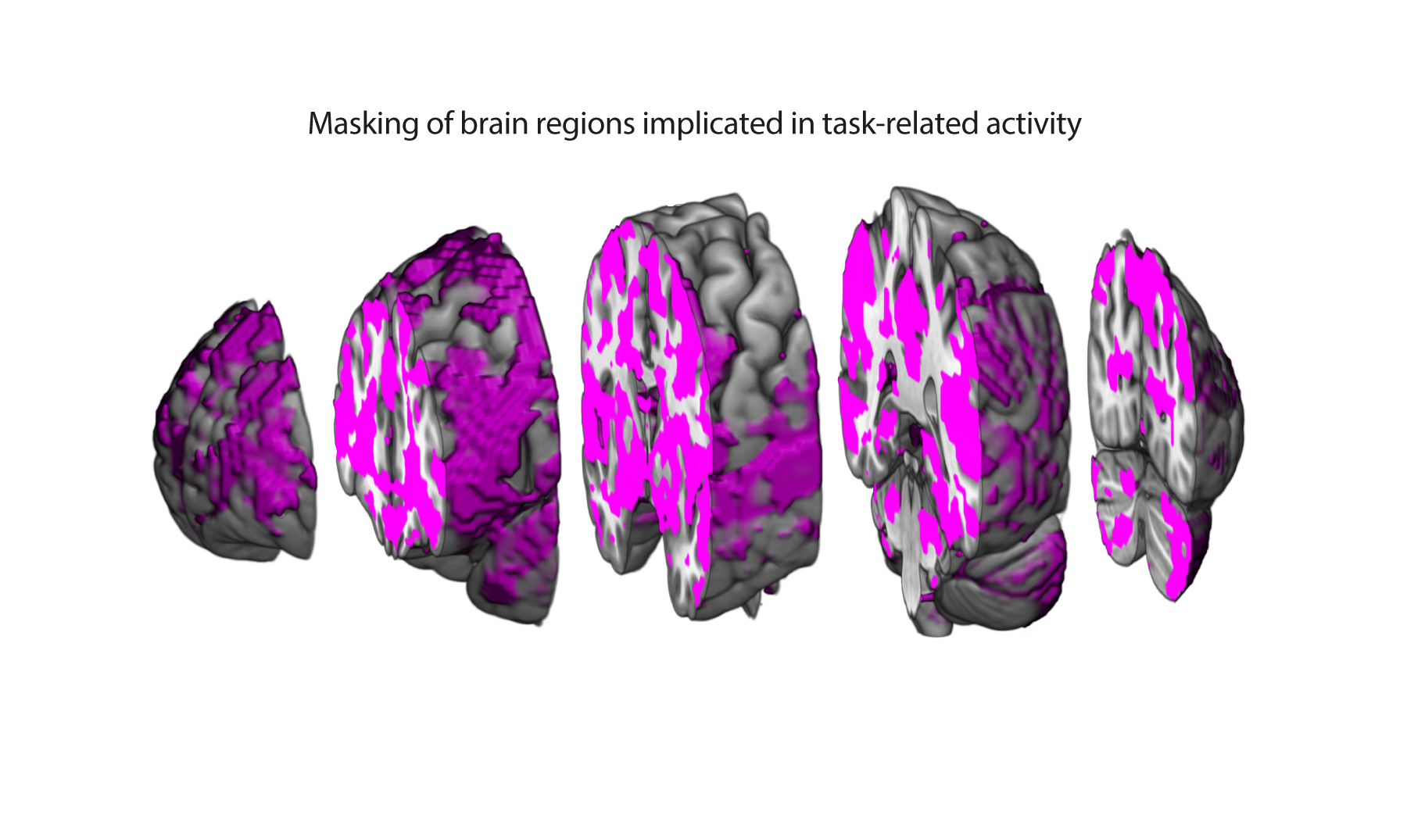

**Supplementary Figure 4.** A brain mask was used to limit the correlational analysis of behavioural and neural responses to those voxels implicated in task-related activity. The mask included all voxels yielding a significant F-contrast across single-participant parameter estimates of all experimental conditions (*p*_uncorrected_ < 0.05).

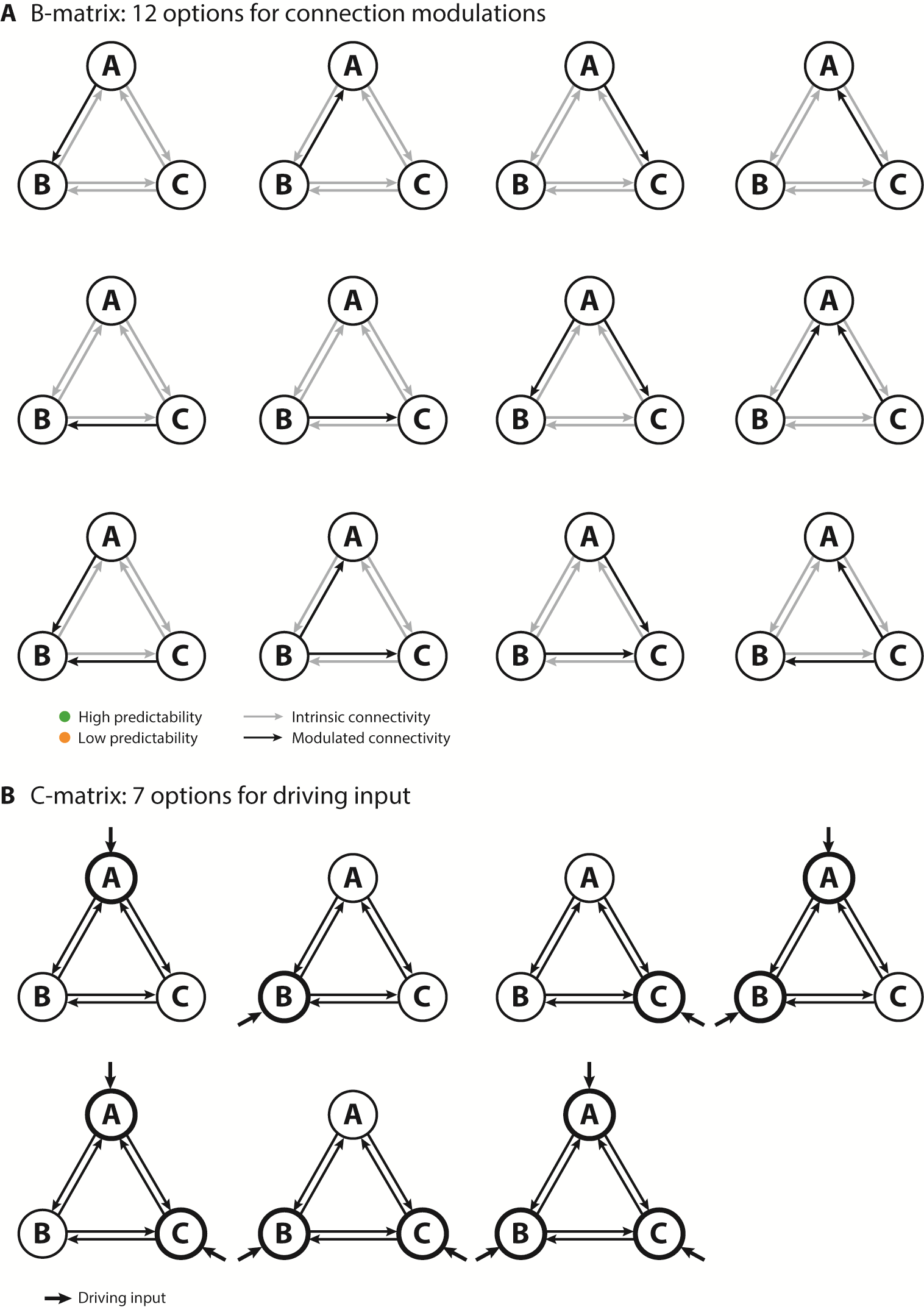

**Supplementary Figure 5.** Schematic overview of the model space for the semantic as well as the cingulo-opercular DCMs. **(A)** Modulation of intrinsic connectivity (arrows) by high (green) and low (orange) predictability. Modulated connections are highlighted in black. **(B)** Fat arrows indicate driving input. A, B and C represent three distinct brain regions.

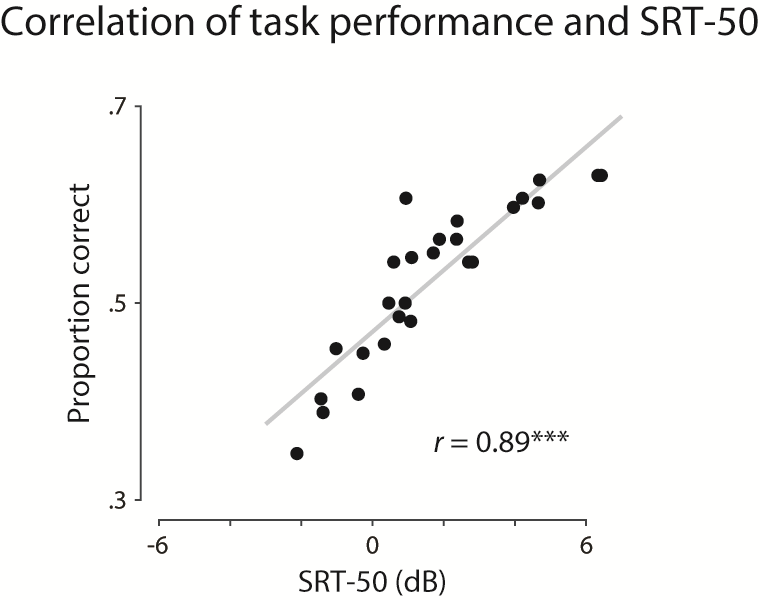

**Supplementary Figure 6.** The SRT-50 determined in the adaptive tracking procedure prior to the experiment was strongly correlated with the proportion of correctly repeated keywords across all conditions (*r* = 0.89, *p* < 0.001, BF_10_ > 1,000).

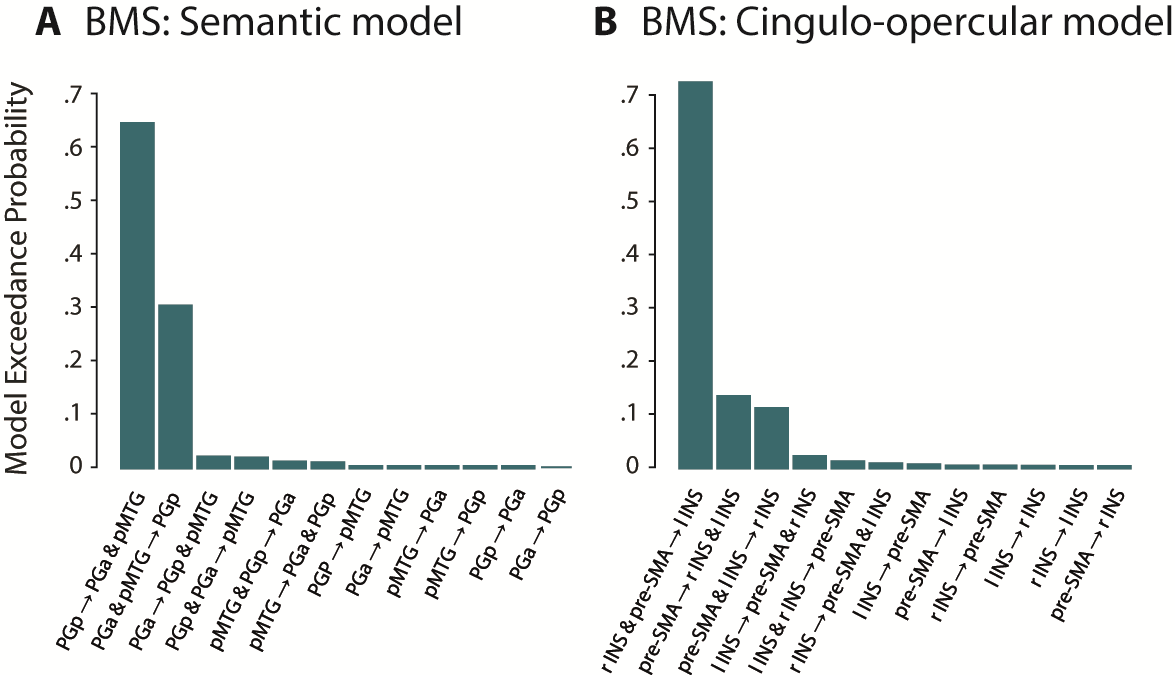

**Supplementary Figure 7.** Exceedance probabilities for all DCM model families estimated using Bayesian Model Selection (BMS). **(A)** Model families within the semantic network DCM. **(B)** Model families within the cingulo-opercular network DCM.

**Supplementary Table 1.** Endogenous and modulatory parameter estimates (Hz) of the semantic network DCM.

| **Connection** | | |  | **Mean** | **SD** | ***t*** | ***p*** | ***d*** | **BF_10_** |
| --- | --- | --- | --- | --- | --- | --- | --- | --- | --- |
| Endogenous Parameters | | |  | | | | | | |
| PGa | → | PGp |  | 0.07 | 0.38 | 1.00 | 0.326 | 0.20 | 0.33 |
| PGa | → | MTG |  | 0.04 | 0.36 | 0.59 | 0.562 | 0.12 | 0.24 |
| PGp | → | PGa |  | 0.04 | 0.31 | 0.69 | 0.499 | 0.14 | 0.28 |
| PGp | → | MTG |  | 0.11 | 0.40 | 1.41 | 0.170 | 0.28 | 0.50 |
| **MTG** | **→** | **PGa** |  | **0.18** | **0.29** | **3.22** | **0.003** | **0.63** | **11.61** |
| **MTG** | **→** | **PGp** |  | **0.20** | **0.29** | **3.55** | **0.002** | **0.70** | **23.13** |
| **PGa** | **↺** |  |  | **–0.18** | **0.11** | **–8.24** | **< 0.001** | **–1.62** | **>100.00** |
| **PGp** | **↺** |  |  | **–0.12** | **0.16** | **–3.69** | **0.001** | **–0.72** | **31.91** |
| **MTG** | **↺** |  |  | **–0.15** | **0.16** | **–4.84** | **< 0.001** | **–0.95** | **>100.00** |
| Modulatory Parameters | | |  | | | | | | |
|  | | | Predictability | Intelligibility | | Mean | SD |  | |
| PGp | → | PGa | High | Low | | **–**0.02 | 1.12 |  |  |
| PGp | → | PGa | High | Medium | | 0.07 | 0.71 |  |  |
| PGp | → | PGa | High | High | | 0.27 | 0.99 |  |  |
| PGp | → | MTG | High | Low | | –0.11 | 0.85 |  |  |
| PGp | → | MTG | High | Medium | | –0.02 | 0.85 |  |  |
| PGp | → | MTG | High | High | | –0.18 | 1.13 |  |  |
| PGp | → | PGa | Low | Low | | –0.37 | 0.93 |  |  |
| PGp | → | PGa | Low | Medium | | –0.33 | 0.83 |  |  |
| PGp | → | PGa | Low | High | | 0.21 | 0.82 |  |  |
| PGp | → | MTG | Low | Low | | –0.24 | 0.94 |  |  |
| PGp | → | MTG | Low | Medium | | –0.89 | 0.71 |  |  |
| PGp | → | MTG | Low | High | | 0.07 | 0.79 |  |  |

Parameters were estimated using Bayesian model averaging (BMA) across models of the winning family and significance was assessed by means of *t*-tests. Significant parameters are highlighted in bold. Effect sizes are reported as Cohen’s *d* and Bayes Factor (BF_10_).

**Supplementary Table 2.** Results of the endogeneous and modulatory parameter estimates (Hz) of the cingulo-opercular DCM.

| **Connection** | | | |  | | **Mean** | **SD** | ***t*** | | ***p*** | ***d*** | **BF10** |
| --- | --- | --- | --- | --- | --- | --- | --- | --- | --- | --- | --- | --- |
| Endogenous Parameters | | |  | | | | | | | | | |
| pre-SMA | → | lIns |  | | –0.04 | | 0.30 | –0.67 | 0.511 | | –0.13 | 0.22 |
| pre-SMA | → | rIns |  | | –0.09 | | 0.24 | –1.84 | 0.078 | | –0.36 | 0.90 |
| **lIns** | **→** | **pre-SMA** | | | **0.29** | | **0.25** | **6.08** | **< .001** | | **1.19** | **>100.00** |
| **lIns** | **→** | **rIns** |  | | **0.25** | | **0.38** | **3.36** | **0.003** | | **0.66** | **15.28** |
| **rIns** | **→** | **pre-SMA** | | | **0.18** | | **0.32** | **2.90** | **0.008** | | **0.57** | **5.96** |
| rIns | → | lIns |  | | 0.09 | | 0.31 | 1.43 | 0.164 | | 0.28 | 0.51 |
| **pre-SMA** | **↺** |  |  | | –**0.15** | | **0.15** | –**4.89** | **< .001** | | –**0.96** | **>100.00** |
| **lIns** | **↺** |  |  | | –**0.16** | | **0.12** | –**6.83** | **< .001** | | –**1.34** | **>100.00** |
| **rIns** | **↺** |  |  | | –**0.09** | | **0.16** | –**2.74** | **0.011** | | –**0.54** | **4.29** |

| Modulatory Parameters | | |  | | | |
| --- | --- | --- | --- | --- | --- | --- |
|  | | | Predictability | Intelligibility | Mean | SD |
| **pre-SMA** | **→** | **lIns** | High | Low | –0.01 | 0.89 |
| **pre-SMA** | **→** | **lIns** | High | Medium | –0.12 | 0.63 |
| **pre-SMA** | **→** | **lIns** | High | High | –0.20 | 1.02 |
| **rIns** | **→** | **lIns** | High | Low | –0.33 | 0.78 |
| **rIns** | → | lIns | High | Medium | –0.06 | 0.78 |
| **rIns** | → | lIns | High | High | –0.28 | 0.77 |
| **pre-SMA** | **→** | **lIns** | Low | Low | –0.42 | 0.77 |
| **pre-SMA** | **→** | **lIns** | Low | Medium | –0.05 | 0.65 |
| **pre-SMA** | **→** | **lIns** | Low | High | –0.02 | 0.56 |
| **rIns** | **→** | **lIns** | Low | Low | 0.09 | 0.92 |
| **rIns** | → | lIns | Low | Medium | –0.11 | 0.77 |
| **rIns** | → | lIns | Low | High | –0.13 | 0.61 |

Parameters were estimated using Bayesian model averaging (BMA) across models of the winning family and significance was assessed by means of *t*-tests. Significant parameters are highlighted in bold. Effect sizes are reported as Cohen’s *d* and Bayes Factor (BF_10_).
